# Pathology of natural infection with highly pathogenic avian influenza virus (H5N1) clade 2.3.4.4b in wild terrestrial mammals in the United States in 2022

**DOI:** 10.1101/2023.03.10.532068

**Authors:** EJ Elsmo, A Wünschmann, KB Beckmen, LB Broughton-Neiswanger, EL Buckles, J Ellis, SD Fitzgerald, R Gerlach, S Hawkins, HS Ip, JS Lankton, EM Lemley, JB Lenoch, ML Killian, K Lantz, L Long, R Maes, M Mainenti, J Melotti, ME Moriarty, S Nakagun, RM Ruden, V Shearn-Bochsler, D Thompson, MK Torchetti, AJ Van Wettere, AG Wise, AL Lim

## Abstract

This article describes the first detections of disease due to natural infection with highly pathogenic avian influenza virus (HPAIv) H5N1 of the Eurasian lineage goose/Guangdong clade 2.3.4.4b in wild terrestrial mammals throughout the United States during 2021-2022. Affected mammalian species include 50 red foxes (*Vulpes vulpes*), 6 striped skunks (*Mephitis mephitis*), 4 raccoons (*Procyon lotor*), 2 bobcats (*Lynx rufus*), 2 Virginia opossums (*Didelphis virginiana*), 1 coyote (*Canis latrans*), 1 fisher (*Pekania pennanti*), and 1 gray fox (*Urocyon cinereoargenteus*). Infected mammals primarily exhibited neurological signs. Necrotizing meningoencephalitis, interstitial pneumonia, and myocardial necrosis were the most common lesions; however, species variations in lesion distribution were observed. Genotype analysis of sequences from 48 animals indicates that these cases represent spillover infections from wild birds.

## Introduction

Since October 2021, outbreaks of highly pathogenic avian influenza (HPAI) H5N1 belonging to Eurasian lineage, clade 2.3.4.4b have been reported throughout European countries (*1*). Highly pathogenic avian influenza virus (HPAIv) with high genetic similarity to Eurasian lineage goose/Guangdong H5 clade 2.3.4.4b was first detected in the United States in December 2021 through wild bird surveillance and has subsequently spread throughout the continental United States in both wild birds and domestic poultry (*2,3,4*).

In addition to causing disease outbreaks in domestic poultry, currently circulating H5N1 HPAIv appears to be persisting in wild bird reservoirs, with multiple reports of spillover into and clinical infection in various mammalian species, including red fox (*Vulpes vulpes*), Eurasian river otter (*Lutra lutra*), Eurasian lynx (*Lynx lynx*), ferret (*Mustela putorius furo*), European polecat (*Mustela putorius*), and stone marten (*Martes foina*) in European countries in 2021 (*1,5*).

This case series reports on the epidemiology and pathology of natural infections with H5N1 HPAIv in terrestrial wild mammals in the United States concurrent with high levels of circulating HPAIv in wild birds and domestic poultry in 2022.

## Materials and Methods

Case inclusion criteria were confirmed positivity for H5N1 HPAIv infection by real time polymerase chain reaction (PCR) or, in three cases, by having consistent microscopic lesions, positive avian influenza virus immunohistochemistry (IHC) results, and being part of a litter in which other individuals were confirmed positive for H5N1 HPAIv by PCR. These cases represent opportunistic sampling of wild mammals that were reported or found by citizens and either submitted to wildlife rehabilitation centers or collected by regional wildlife professionals and state agency personnel. Clinical observations were variably recorded by citizens, wildlife professionals, rehabilitators, or veterinary care professionals. Antemortem nasal and oropharyngeal swabs were obtained from two red foxes that remained alive and were released to the wild. For the other mammals, partial or complete postmortem examinations were performed by veterinarians or pathologists at regional or federal diagnostic laboratories or wildlife agencies. A variety of samples were collected postmortem, including nasal, oropharyngeal, tracheal, intestinal, rectal, and tissue swabs in viral transport media; tissue samples stored refrigerated or frozen were fixed in 10% neutral buffered formalin (Supplementary Table 4).

The standardized protocols for the National Animal Health Laboratory Network (NAHLN) were utilized for HPAIv real-time PCR testing on a variety of sample types including swabs and tissues. Total nucleic acids were extracted from fresh samples using the KingFisher Flex or KingFisher Purification System platforms and MagMAX-96 Viral RNA Isolation Kit or MagMAX Pathogen RNA/DNA Kit (Life Technologies, Carlsbad, California), following the manufacturer’s protocol. A general influenza A virus (IAV) PCR targeting the conserved region of the IAV matrix gene was performed (*6,7*). Further avian influenza subtyping was performed using two IAV H5 subtyping assays: the avian influenza H5 subtype PCR targeting the hemagglutinin gene for the North American, Eurasian, and Mexican lineage of avian influenza (*6,7,8,9*), and an H5 2.3.4.4b-specific PCR developed in collaboration with USDA Southeast Poultry Research Laboratory (SEPRL; Real-Time RT-PCR Assay for the Detection of Goose/Guangdong lineage Influenza A subtype H5, clade 2.3.4.4; NVSL-WI-1732).

Samples or nucleic acid extracts with non-negative influenza A results were shipped frozen to the National Veterinary Services Laboratories (NVSL) in Ames, Iowa for confirmation and characterization. Testing at NVSL included an H5 clade 2.3.4.4 pathotyping assay and an assay targeting N1 for rapid pathotyping (SEPRL; Real-Time RT-PCR Assay for Pathotyping Goose/Guangdong lineage Influenza A subtype H5, clade 2.3.4.4; NVSL-WI-1767) and neuraminidase subtyping (SEPRL; Real-Time RT-PCR Assay for the Detection of Eurasian-lineage Influenza A Subtype N1; NVSL-WI-1768). Influenza A viruses were sequenced directly from samples as previously described *(2)*. RAxML was used to generate phylogenetic trees, and tables of single nucleotide polymorphisms (SNPs) were created using the vSNP pipeline (https://github.com/USDA-VS/vSNP) with a reference composed of six segments from an H5N1 2.3.4.4b clade virus and two segments from North American origin wild bird viruses (see Supplementary Table 1).

In some cases, extracted total nucleic acids were also screened for other viral etiologies by established real-time PCR assays. Fresh brain tissue was collected from a subset of cases and submitted for rabies virus fluorescent antibody or real-time PCR testing. Fresh samples of pooled liver, kidney, spleen, and a swab of the abdominal cavity of one red fox were submitted for routine aerobic culture. A sample of liver from the gray fox (*Urocyon cinereoargenteus*) was screened for anticoagulant rodenticides by liquid chromatography/mass spectrometry. Formalin-fixed tissues were routinely processed for histopathology and were evaluated by veterinary pathologists at multiple institutions. A subset of tissues from some animals were also processed for immunohistochemistry (IHC) for IAV antigen using a monoclonal antibody to influenza A viral nucleoprotein (NP) and/or canine distemper virus antigen using a monoclonal antibody to canine distemper virus NP. All ancillary testing procedures were performed according to validated procedures at federal and state diagnostic laboratories accredited by the American Association of Veterinary Laboratory Diagnosticians.

## Results

Between April 1 and July 21, 2022, HPAIv was detected in 67 wild mammals from 10 states in the continental United States: Alaska, Idaho, Iowa, Michigan, Minnesota, New York, North Dakota, Utah, Washington, and Wisconsin, in 44 different counties (Fig 1 A-C). Species included 50 red foxes (*Vulpes vulpes*), 6 striped skunks (*Mephitis mephitis*), 4 raccoons (*Procyon lotor*), 2 bobcats (*Lynx rufus*), 2 Virginia opossums (*Didelphis virginiana*), 1 coyote (*Canis latrans*), 1 fisher (*Pekania pennanti*), and 1 grey fox (*Urocyon cinereoargenteus*) (Supplementary Table 2).

**Figure 1.**
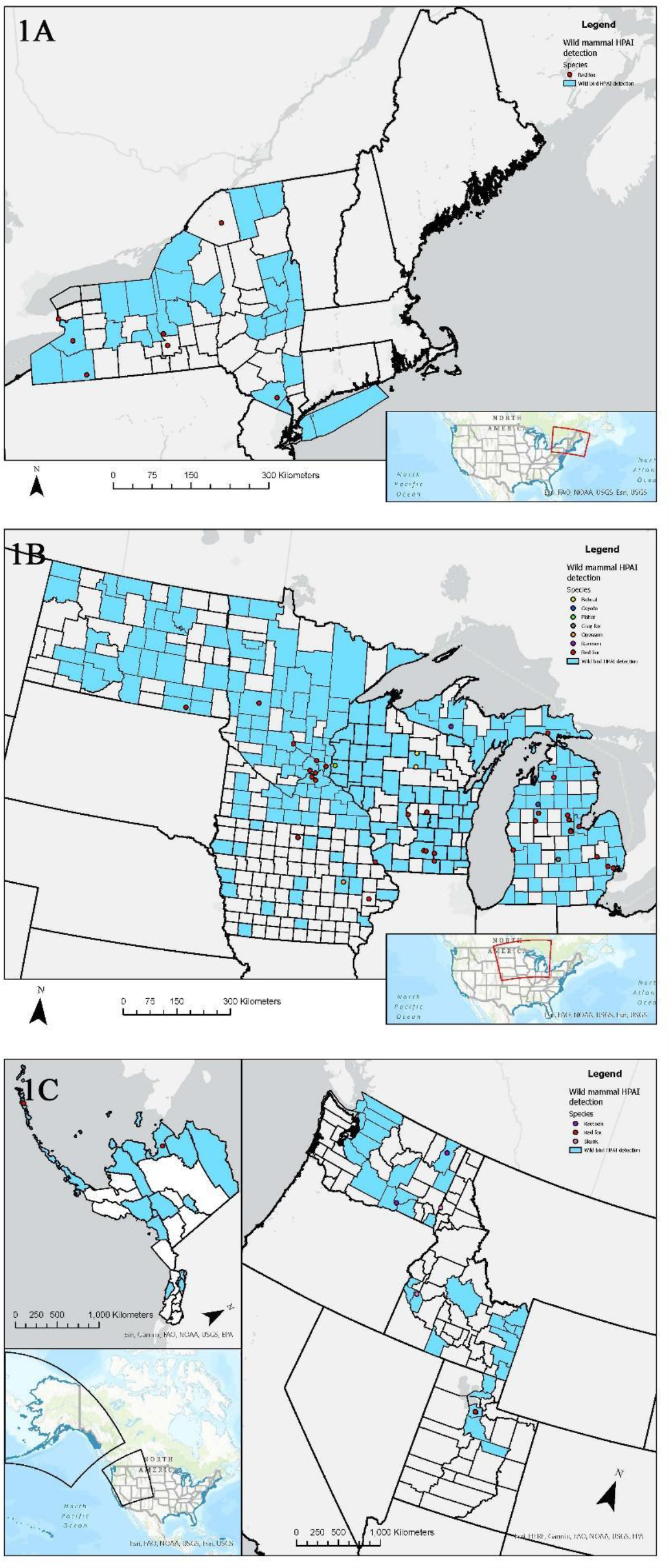
**A-C:** Map of the locations of wild mammals with highly pathogenic avian influenza H5N1 in the United States (A – Eastern, B – Midwestern, C – Western) from April - June 2022 (species depicted by colored points). If exact location could not be identified, the county centroid was used. Counties with positive detections of highly pathogenic avian influenza H5N1 in wild birds from March - August 2022 are indicated in blue.

All red foxes, striped skunks, opossums, 3 of 4 raccoons, and the single coyote were juveniles. A single raccoon, both bobcats, the fisher, and grey fox were adults. Age determinations were based on individual body weight and dentition using published reference ranges (*10*). Within species with greater than 4 individuals where sex was recorded, no sex predilections were apparent (Supplementary Table 2).

For one raccoon, the mechanism of death (natural or euthanasia) was not recorded. Ten animals were found dead. Of the animals found alive, 13 were euthanized in the field and 43 animals were under the care of a wildlife rehabilitator or veterinarian for some duration of time (see Supplementary Table 2). Of these, two red foxes remained alive and clinically normal prior to release, 12 died, and the remainder were euthanized shortly after admission or as a result of poor prognosis.

Among the 57 animals that were found or observed alive, most (n=53) had neurological abnormalities comprising seizures (n=28), ataxia (n=23), tremors (n=16), inappropriate or lack of fear of humans (n=7), vocalization (n=5), circling (n=4), blindness (n=3), torticollis (n=2), nystagmus (n=1), and grimace (n=1) (Supplementary Table 3). Less frequently recorded signs included lethargy (n=28), fever (n=7), diarrhea (n=2), unconsciousness (n=2), recumbence (n=1), paralysis (n=1), and vomiting (n=1). Dyspnea was reported in 2 of 5 skunks, 1 bobcat, and 1 red fox, but was not observed in other species. One red fox (red fox 24) was found as a presumptively abandoned juvenile, had no reported clinical signs, but was thin and dehydrated on admission.

At postmortem evaluation, gross observations were recorded in 58 animals, 6 of which had evidence of trauma due to either vehicular collision or method of euthanasia. Postmortem conditions varied from fresh to significantly autolyzed and many animals were frozen prior to examination. The majority of animals were in fair to good nutritional condition (n=39). The most consistently observed gross lesions were in the lungs (n=49). The most common pulmonary lesions included congestion (n=42), edema (n=22), failure to collapse (n=18), hemorrhage (n=18), and pleural effusion (n=6) (Fig. 2A). The most common brain lesion was hemorrhage (n=11) and congestion (n=7) with hydrocephalus and malacia reported in one red fox each. Other lesions included pallor (n=8), congestion (n=7), enlargement (n=6), and hemorrhage (n=1) in the liver, congestion (n=4) and cortical hemorrhage (n=1) in the kidney. Pericardial effusion (n=3), petechiae (n=2), and myocardial pallor (n=2) were infrequently noted in the heart (Fig. 2B). In the gastrointestinal tract, nematode parasitism was relatively common (n=15), but other rarely observed lesions included congestion, hemorrhage, and loose feces. Three red foxes had gastric contents that included feathers (Fig. 2C). Three red foxes had mild ocular discharge. One grey fox had severe hemorrhage in all body cavities.

**Figure 2.**
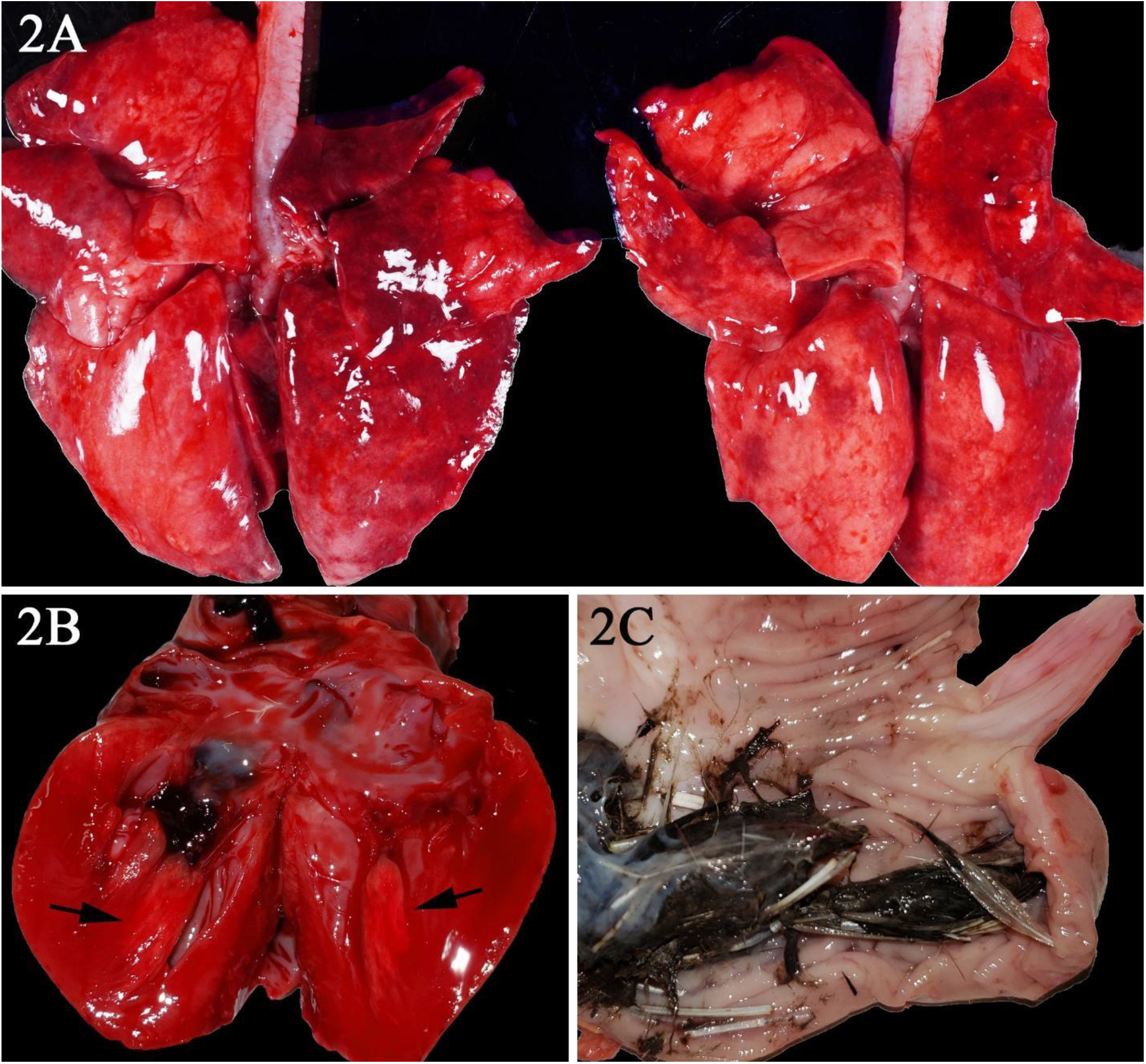
**A-C**. Gross photographs of postmortem lesions from red foxes naturally infected with highly pathogenic avian influenza virus. A – The lungs have failed to collapse and are diffusely edematous and mottled pink to dark red. B – Cross section of the left ventricle of the heart revealing a focal region of myocardial pallor in the papillary muscle (arrows). C – The stomach contents with feathers.

Histopathological findings were recorded in 55 animals (summarized in Supplementary Table 4). Of the 54 animals with at least one section of brain suitable for histopathological evaluation, all but two skunks and one grey fox had brain lesions. Although the brain regions evaluated were not consistent amongst all animals, the most frequently affected region reported was the cerebral cortex (n=46), followed by the brainstem (n=22), thalamus (n=20), hippocampus (n=17), frontal lobe (n=16), and cerebellum (n=12). Brain lesions had a multifocal and random distribution, affected both gray and white matter, and sometimes exhibited a periventricular distribution. Lesions primarily consisted of regions of malacia and inflammation (Fig. 3A). Acidophilic neuronal necrosis was prominent and often associated with satellitosis (Fig. 3B) or karyorrhectic debris (Fig. 3C). Laminar neuronal necrosis within the hippocampus was evident in some cases (Fig. 3D). Glial nodules and reactive astrocytes were common in affected regions. Infiltrating the leptomeninges, Virchow-Robin spaces, and often extending into the parenchyma were low to moderate numbers of predominantly lymphocytes and plasma cells (Fig. 3A and 3B). Histiocytic, neutrophilic, and eosinophilic infiltrates were noted in some cases; and in some presumptively subacute lesions reactive histiocytes predominated. Fibrinoid vascular necrosis was uncommonly reported but was marked in one red fox that was co-infected with canine adenovirus. Notably, histopathological lesions in the brain were absent to mild in all skunks evaluated. Although autolysis may have obscured subtle lesions in the two opossum joeys, brain lesions were mainly affecting the cerebral meninges.

**Figure 3.**
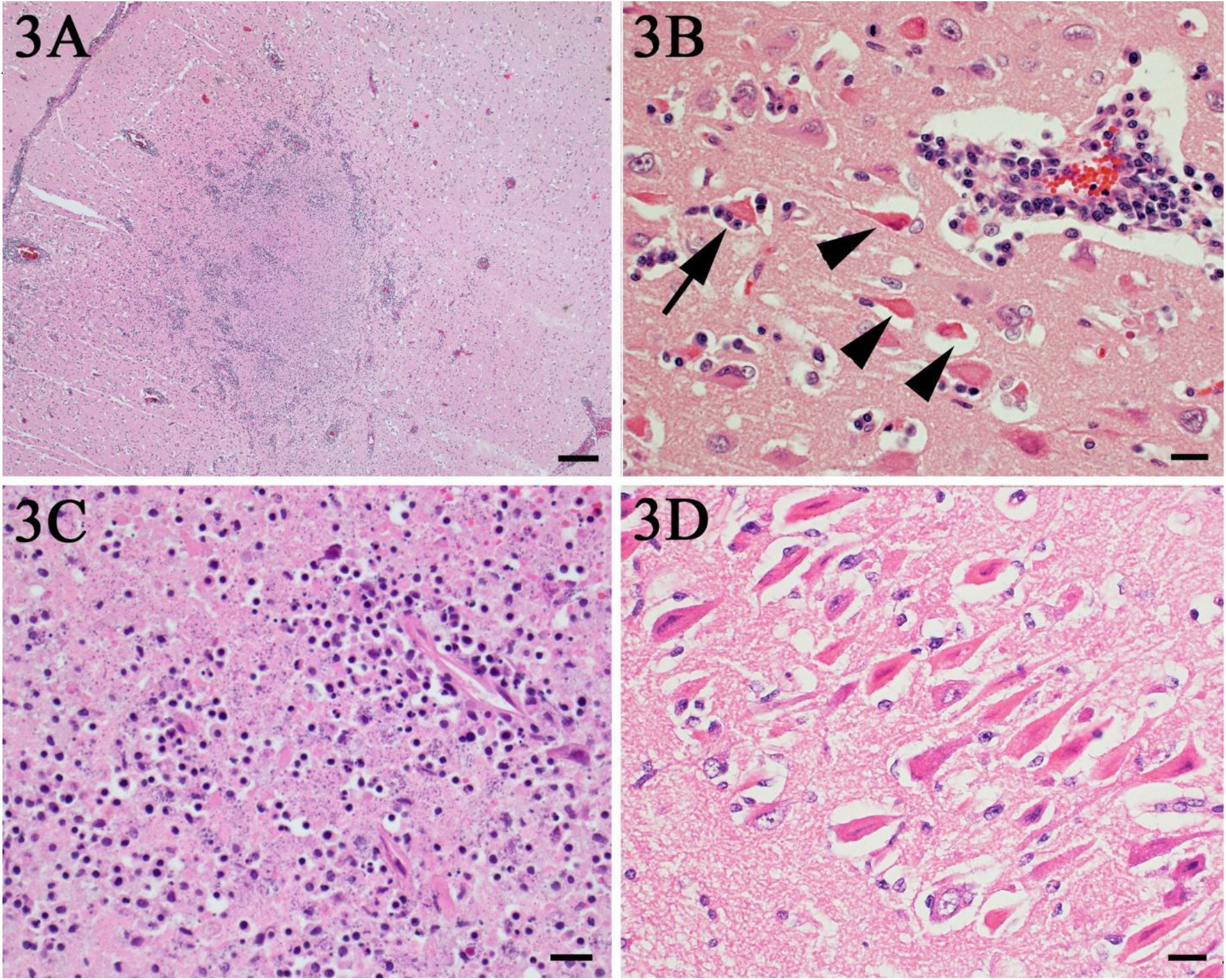
**A-H**. Photomicrographs of histopathological lesions in red foxes naturally infected with highly pathogenic avian influenza. A – Throughout the brain there are multifocal regions of necrosis and hypercellularity. H&E stain. Bar = 200μm. B – Within the grey matter there is prominent neuronal necrosis (arrow heads), satellitosis (arrow), and reactive astrocytes. A vessel is surrounded by lymphocytes and plasma cells. H&E stain. Bar = 20 μm. C – In areas of necrosis within the brain, there is often abundant, stippled, basophilic karyorrhectic debris. H&E stain. Bar = 20 μm. D - Within the hippocampus, there are numerous shrunken, angular, and acidophilic (necrotic) neurons in a laminar pattern. H&E stain. Bar = 20 μm.. E – Within the lung, there is diffuse vascular congestion. Alveoli contain fibrin, hemorrhage, and edema fluid. H&E stain. Bar = 50 μm. F – Regions of cardiomyocyte necrosis in the heart are often mineralized. H&E stain. Bar = 20 μm. G – Within the brain, there is positive nuclear and cytoplasmic staining of neuron cell bodies and processes. Avian influenza virus monoclonal immunohistochemistry. Bar = 20 μm. H – There is scattered positive nuclear and cytoplasmic staining of bronchiolar epithelia l cells and interstitial macrophages in the lung. Monoclonal immunohistochemistry to influenza A virus nucleoprotein. Bar = 20 μm.

**Figure 3.**
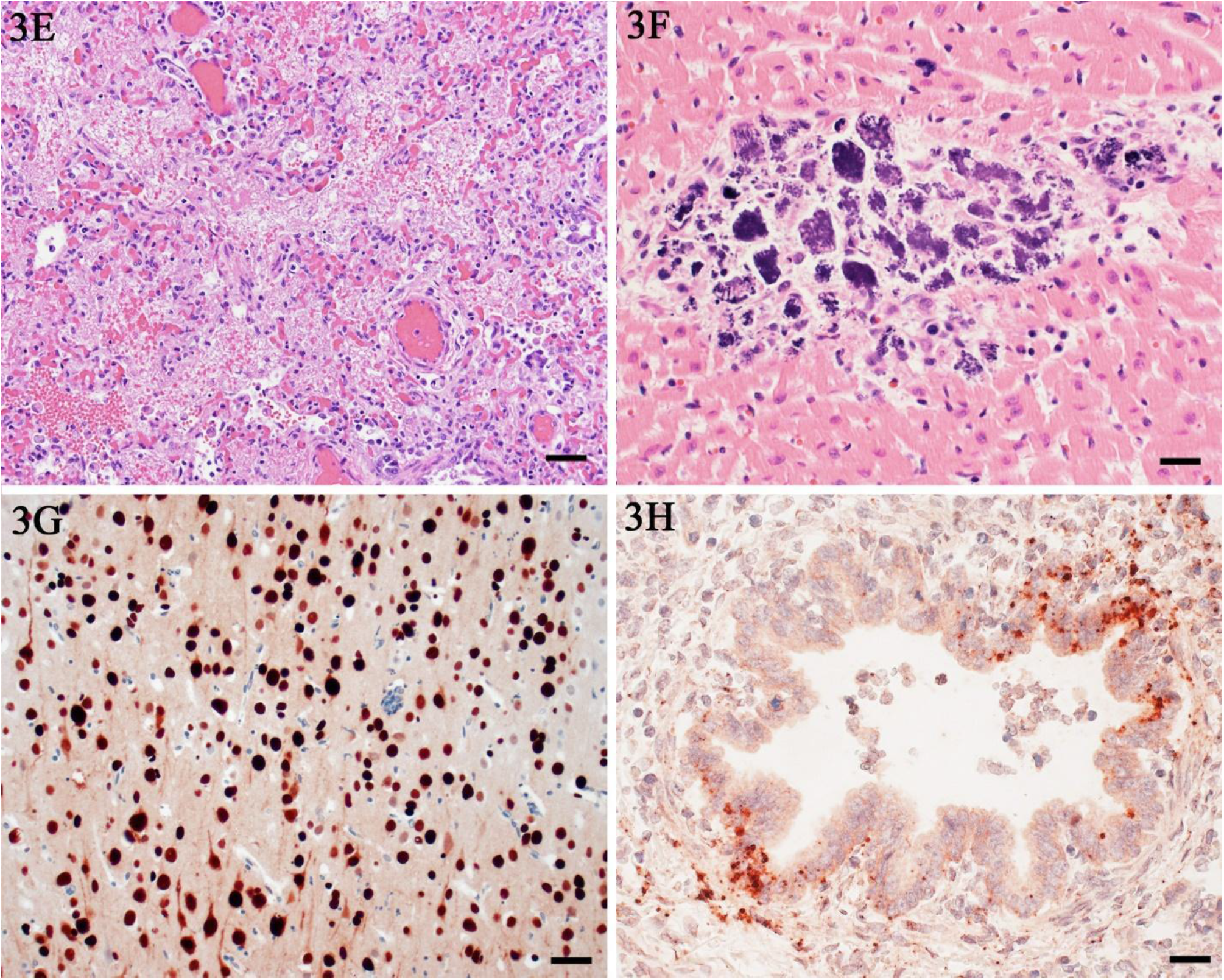
E – Within the lung, there is diffuse vascular congestion. Alveoli contain fibrin, hemorrhage, and edema fluid. H&E stain. Bar = 50 μm. F – Regions of cardiomyocyte necrosis in the heart are often mineralized. H&E stain. Bar = 20 μm. G – Within the brain, there is positive nuclear and cytoplasmic staining of neuron cell bodies and processes. Avian influenza virus monoclonal immunohistochemistry. Bar = 20 μm. H – There is scattered positive nuclear and cytoplasmic staining of bronchiolar epithelia l cells and interstitial macrophages in the lung. Monoclonal immunohistochemistry to influenza A virus nucleoprotein. Bar = 20 μm.

Multifocal necrotizing interstitial pneumonia was the second most commonly reported microscopic lesion, affecting 47 animals. Alveoli contained variable amounts of fibrin, edema fluid, hemorrhage, alveolar macrophages, and occasionally neutrophils (Fig. 3E). Alveolar septa were sometimes thickened by neutrophils, macrophages, perivascular lymphocytes and plasma cells, and aggregates of fibrin. Type II pneumocyte hyperplasia was rarely observed, and few animals had necrotizing, neutrophilic, or histiocytic bronchopneumonia. The majority of lung lesions were acute, excepting one fox (red fox 30) and one bobcat (bobcat 1). Concurrent lungworm parasitism was noted in three raccoons, one bobcat, and the fisher.

Multifocal myocardial necrosis, most commonly affecting the left ventricular wall, was present in roughly half of the animals (n=29). Mineralization of affected cardiomyocytes was often noted (Fig. 3F). Mild lymphoplasmacytic or histiocytic infiltrates and regions of fibrosis were infrequently reported in conjunction with regions of myocardial necrosis. No heart lesions were observed in the striped skunks, the opossum joeys, the fisher, or the grey fox.

Randomly distributed foci of acute hepatic necrosis were observed in 22 animals and notably, in all five striped skunks. Necrosis varied from liquefactive to coagulative, and dystrophic mineralization of hepatocytes was reported in some cases. Mixed neutrophilic, histiocytic, and lymphoplasmacytic inflammation was occasionally observed associated with regions of necrosis. Chronic cholangitis with cholestasis was noted in one bobcat.

Lymphoid depletion in the spleen, thymus, and lymph nodes was observed in 28 animals and was most consistent in the striped skunks and raccoons. Lymphoid necrosis was prominent in the spleen (n=5), lymph nodes (n=4), and Peyer’s patches of the ileum (n=1) of the striped skunks, and in the thymus and intestinal Peyer’s patches of one raccoon.

Rarely, foci of inflammation or necrosis were observed in the kidney, tongue, and gastrointestinal tract in the red fox kits. A focal area of acute pancreatic necrosis and multifocal lymphoplasmacytic pancreatitis were noted in one red fox each. With the exception of a few small clusters of lymphocytes and plasma cells within the photoreceptor layer of the retina of one red fox, no other histopathologic lesions were reported in any of the examined eyes from other red foxes, including the three red foxes reported as clinically blind. The single grey fox had no microscopic lesions.

Immunohistochemical analysis (IHC) for avian influenza antigen was performed on a variety of tissues in 29 animals, including brain tissue from 17 red foxes, 2 bobcats, 2 opossums, and 1 raccoon. Of the 13 red foxes that had any immunoreactivity in the brain, positively staining neuronal cell bodies and processes were detected in the cerebral cortex (n=13), thalamus (n=6), hippocampus (n=3), and brainstem (n=1) (Figure 3G). Both opossums had immunoreactivity in the cortex, and one bobcat had immunoreactivity in the cortex, thalamus, and hippocampus. The other bobcat, which had subacute brain lesions primarily populated by reactive histiocytes, exhibited no immunoreactivity in the brain. The single raccoon had immunoreactivity within the brainstem, but immunoreactivity within the cerebral cortex was equivocal.

Immunoreactivity within lung tissue was limited to scattered mild staining of bronchial gland epithelial cells and interstitial macrophages (Figure 3H) in 2 of 15 red foxes that had IHC performed on lung. Two striped skunks and one raccoon had similar scattered immunoreactivity within the lung. One of two opossums had strong multifocal nuclear and cytoplasmic immunoreactivity within pneumocytes and to a lesser extent alveolar macrophages and bronchial and bronchiolar epithelium. Neither bobcat had immunoreactivity in the lung.

In the five red foxes and one raccoon that had IHC performed on heart tissue, three red foxes and the single raccoon exhibited immunoreactivity of scattered cardiac myofibers and interstitial macrophages surrounding foci of necrosis.

All five striped skunks that had IHC performed in the liver had immunoreactivity, primarily within hepatocytes surrounding necrotic foci whereas only one of three red foxes tested had immunoreactivity in liver lesions. Immunoreactivity was not detected in the spleens of three red foxes and one raccoon tested. However, there was scattered immunoreactivity in lymphoid organs, including within regions of thymic lymphoid necrosis in a raccoon and in the peripancreatic lymph node of a skunk. A single red fox had immunoreactivity in a few mesenteric arterioles.

A variety of sample types from 64 animals were tested by PCR in NAHLN laboratories using several assays that can detect IAV. With the exception of one red fox (assay performed on formalin-fixed paraffin embedded brain tissue at the New York Animal Health and Disease Center), all animals with non-negative PCR results had at least one sample test positive at NVSL. The majority of samples were uniformly positive using the general IAV, H5 subtype, and H5 clade 2.3.4.4b subtype PCR assays, when performed (summarized in Supplementary Table 5). Of 35 animals that had 2 or more sample types tested, brain samples most frequently had the strongest amplification signal (n=21). In 3 red foxes, swabs from the nasal and/or oropharyngeal cavities had the strongest amplification signal, and in 2 red foxes the lung had the strongest signal. The nasal or oropharyngeal swab was the only positive sample type in two red foxes (red fox 19 and 24), and a pooled nasal and oral swab was the only positive sample type in the grey fox. In 16 animals with nasal and oropharyngeal swabs, oropharyngeal swabs frequently had the stronger amplification signal (n=10).

Full genome sequence data were obtained directly from 77 samples representing 48 individual animals across seven different species (bobcat (n=1), coyote (n=1), fisher (n=1), opossum (n=2), raccoon (n=2), red fox (n=37) and skunk (n=4)), in 10 states (Alaska (AK), Iowa (IA), Idaho (ID), Michigan (MI), Minnesota (MN), North Dakota (ND), New York (NY), Utah (UT), Washington (WA), Wisconsin (WI)). Among the 48 individual animals, 9 different H5N1 genotypes were identified based on phylogenetic analysis and according to the specific combinations of Eurasian and North American gene segments (Table 3, Table 5). All but one of these genotypes represent reassortments of the Newfoundland-like H5N1 2.3.4.4b virus initially introduced into the Americas via the Atlantic flyway with North American wild bird origin influenza viruses (all have at least North American PB1 and NP genes). The single unreassorted virus is from a red fox in Alaska and represents a separate introduction of the H5N1 2.3.4.4b into the Americas (a Japan-like virus likely introduced via the Bering Strait), based on phylogenetic analysis and high sequence identity to Asian-origin H5N1 viruses across all eight segments (*11*). E627K substitution previously reported to be associated with mammalian adaptation (*12*) was identified in three animals (WA raccoon 2 genotype B2, IA red fox 31 genotype B1.2, and MI red fox 45 genotype B3.2), each from different states and of different genotypes – of note, these were also the most frequently identified genotypes. Analysis of all sequences indicates infections resulted from regional spillovers from wild birds (refer to examples of a subset of genotype B1.2 in Fig. 4A-B and to phylogenetic trees of the two most common genotypes in Fig. 5A-B).

**Figure 4A.**
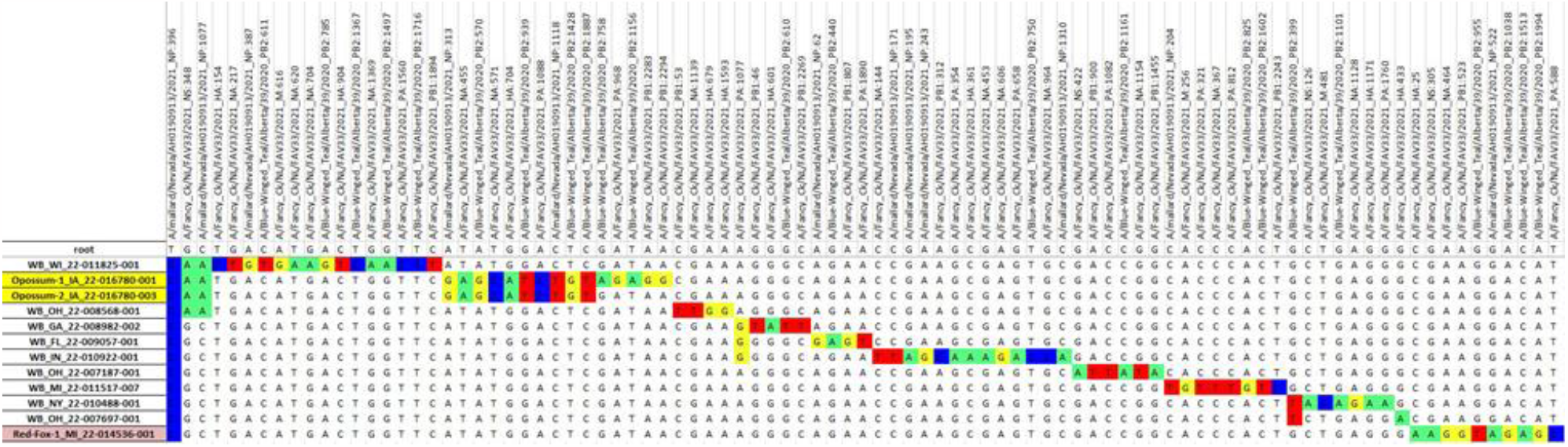
Example of a subset of B1.2 vSNP analysis using whole genome data for H5N1 clade 2.3.4.4b highly pathogenic avian influenza in mammals and wild birds. Data from three samples from one red fox and two samples from one Virginia opossum are highlighted in red and yellow, respectively. Each column represents the nucleotide position relative to the reference virus.

**Figure 4B.**
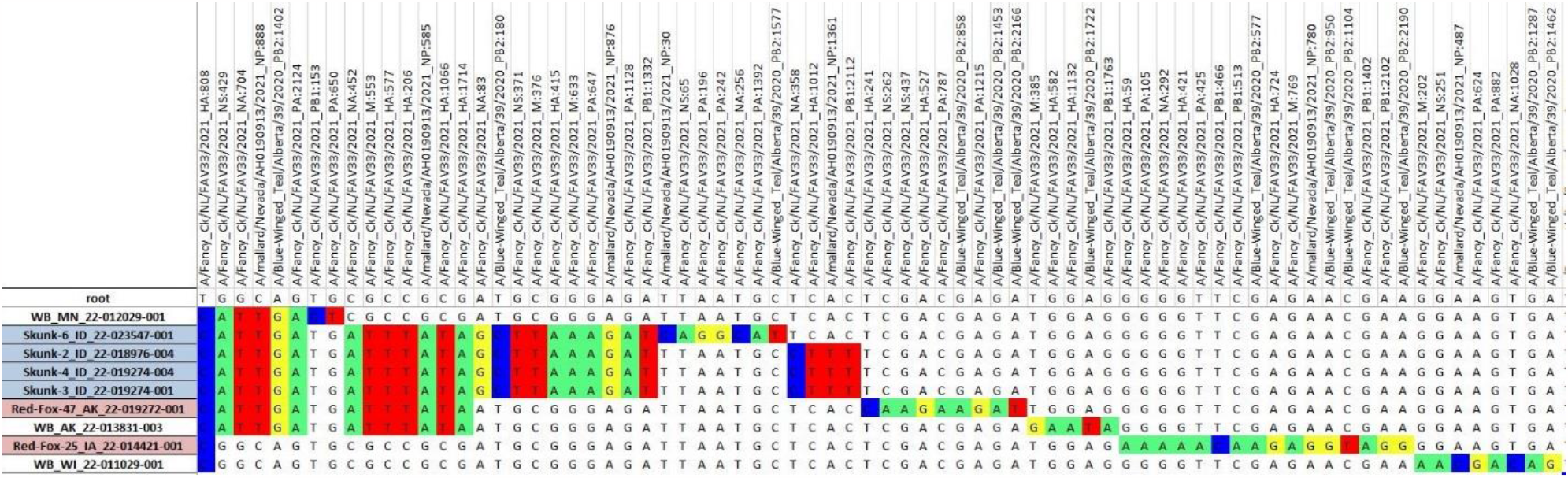
Example of a subset of B3.2 vSNP analysis using whole genome data for H5N1 clade 2.3.4.4b highly pathogenic avian influenza in mammals and wild birds. Data from four samples from four skunks and three samples from two red foxes are highlighted in blue and red, respectively. Each column represents the nucleotide position relative to the reference virus.

**Figure 5 A.**
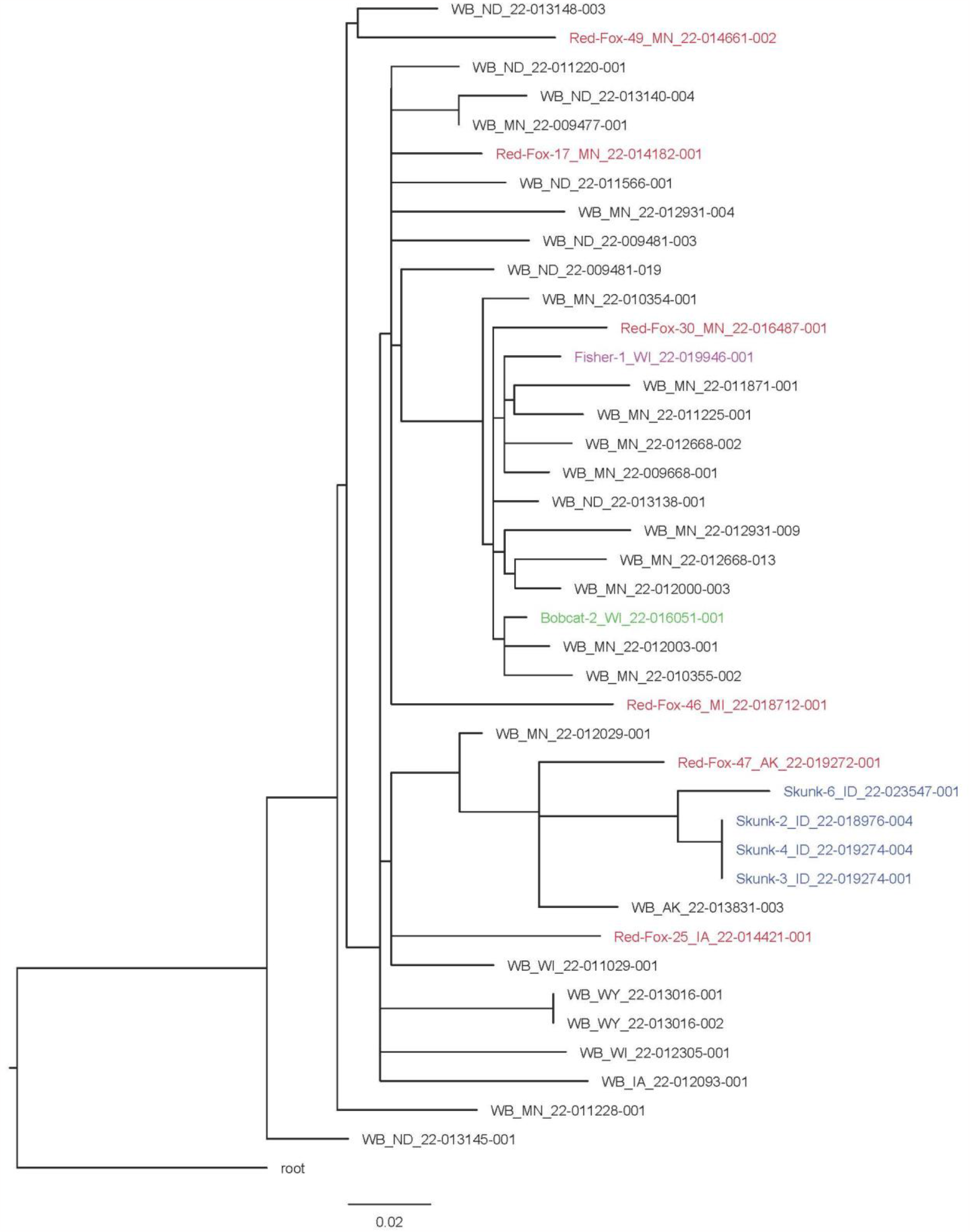
Genotype B3.2 vSNP phylogenetic tree for H5N1 clade 2.3.4.4b highly pathogenic avian influenza in mammals and wild birds. Data from red fox are shown in red, fisher in purple, bobcat in green, and skunk in blue. WB = wild bird. Tree is rooted to the reference sequence A/Fancy_Ck/NL/FAV33/2021.

**Figure 5 B.**
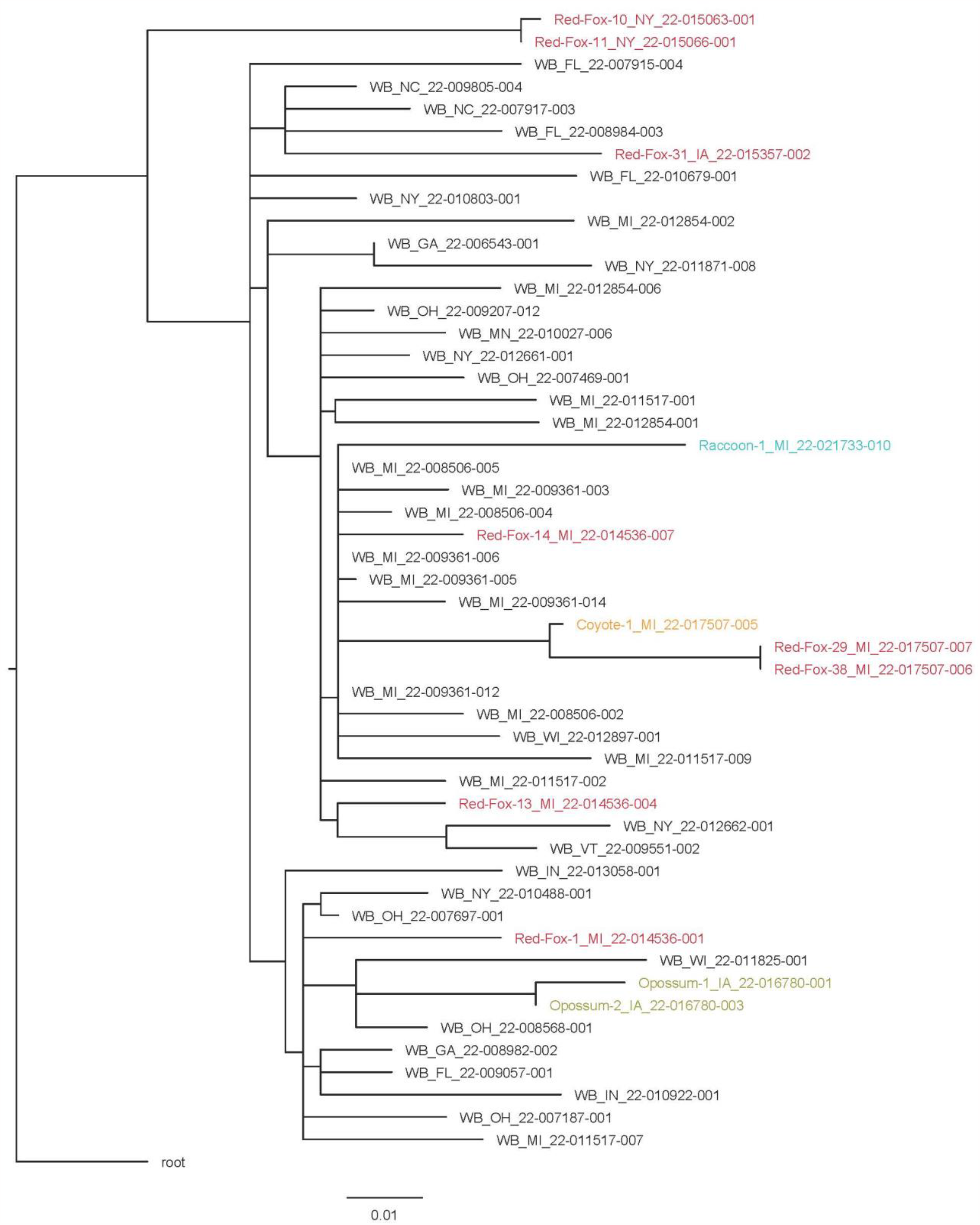
Genotype B1.2 vSNP phylogenetic tree for H5N1 clade 2.3.4.4b highly pathogenic avian influenza in mammals and wild birds. Data from red fox are shown in red, raccoon in teal, coyote in orange, and Virginia opossum in gold. WB = wild bird. Tree is rooted to the reference sequence A/Fancy_Ck/NL/FAV33/2021.

**Table 1.**
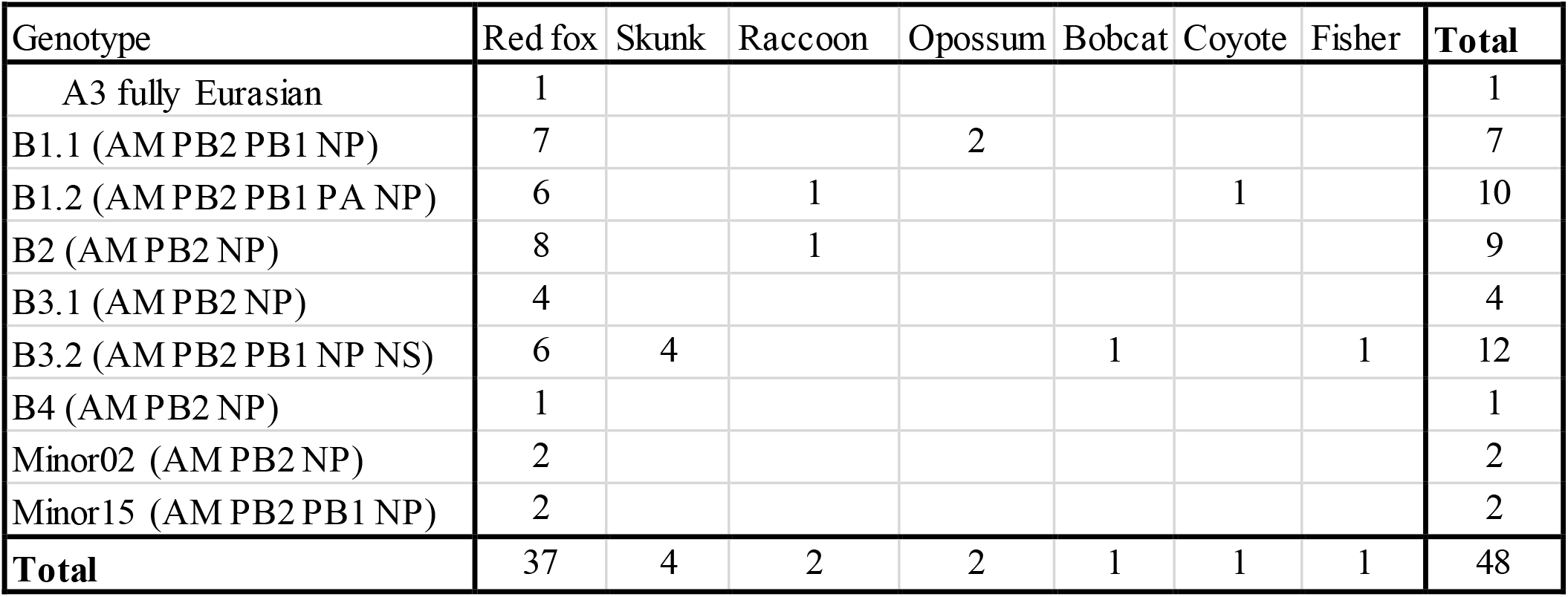
Nine different genotypes were identified from the whole genome sequence data of H5N1 clade 2.3.4.4b highly pathogenic avian influenza from 48 individual animals across seven different species and 10 states (AK, IA, ID, MI, MN, ND, NY, UT, WA, WI); (North American wild bird influenza A segments denoted by AM)

| Genotype | Red fox | Skunk | Raccoon | Opossum | Bobcat | Coyote | Fisher | Total |
| --- | --- | --- | --- | --- | --- | --- | --- | --- |
| A3 fully Eurasian | 1 |  |  |  |  |  |  | 1 |
| B1.1 (AM PB2 PB1 NP) | 7 |  |  | 2 |  |  |  | 7 |
| B1.2 (AM PB2 PB1 PA NP) | 6 |  | 1 |  |  | 1 |  | 10 |
| B2 (AM PB2 NP) | 8 |  | 1 |  |  |  |  | 9 |
| B3.1 (AM PB2 NP) | 4 |  |  |  |  |  |  | 4 |
| B3.2 (AM PB2 PB1 NP NS) | 6 | 4 |  |  | 1 |  | 1 | 12 |
| B4 (AM PB2 NP) | 1 |  |  |  |  |  |  | 1 |
| Minor02 (AM PB2 NP) | 2 |  |  |  |  |  |  | 2 |
| Minor15 (AM PB2 PB1 NP) | 2 |  |  |  |  |  |  | 2 |
| **Total** | 37 | 4 | 2 | 2 | 1 | 1 | 1 | 48 |

Ancillary testing for other disease agents was performed on many of these animals (summarized in Supplementary Table 6). One of 36 red foxes tested for rabies virus was positive. Canine distemper virus was not detected in 31 animals from which samples were tested. One red fox (red fox 19) was concurrently infected with canine adenovirus, as confirmed by PCR testing of brain tissue and consistent microscopic findings in the brain. One red fox (red fox 12) was diagnosed with concurrent *Escherichia coli* enteritis and septicemia based on histopathological findings and bacterial culture results. Parvovirus was detected in both bobcats by PCR on samples of brain and lung, and although microscopic evidence of lymphoid depletion was found, lesions consistent with parvoviral enteritis were not identified in either bobcat. No anticoagulant rodenticides were detected in a liver sample from the grey fox.

## Discussion

These cases represent the first detections of the currently circulating highly pathogenic avian influenza virus Eurasian lineage H5 clade 2.3.4.4b in wild terrestrial mammals in the United States. Our findings build on prior reports of natural infections with this HPAIv strain in wild red foxes from the Netherlands in 2021 (*13*) and add to the global list of mammalian species susceptible to H5N1 HPAIv (*14*). Additionally, these cases represent the first reports of natural H5N1 HPAIv infection in wild bobcats, coyotes, grey foxes, fishers, raccoons, striped skunks, and Virginia opossums.

Given the broad scope and ongoing nature of the outbreak, these cases likely represent only a small percentage of the total number and species of mammals infected with the currently circulating strain of HPAIv in the United States. Our report is limited by several factors, including the number of organizations participating in this study, sampling effort at those institutions and their submitting agencies, and the visibility of ill and deceased wildlife on the landscape. Red foxes are the most represented species in this report. Several intrinsic factors related to exposure and infection risk could explain this finding, such as scavenging behavior and dietary preferences, likelihood of sharing environments with infected birds, abundance of immunologically naive animals present during the onset of the avian outbreak, and potentially increased susceptibility to infection in this species. Many of the red foxes in this study were found in urban or peri-urban environments, and extrinsic factors such as human interest in these highly visible animals living near populous areas may have led to increased public reporting. Raccoons, skunks, opossums, and coyotes are also generalist mesopredators frequently encountered in urban and peri-urban areas (*15)*, and reasons why these species were less represented in this study are unclear. Although serological evidence of exposure to avian influenza viruses has been documented in a variety of wild mammal species, few experimental trials have investigated the susceptibility of wild mammals to avian influenza viruses, and even fewer specifically to H5N1 HPAIv (*16*). Further studies on the susceptibility of mammalian species to infection with the currently circulating strains of H5N1 HPAIv may be warranted, especially in light of the unprecedented reassortment of the Newfoundland-like virus with North American wild bird origin influenza viruses (*2*).

The mammals in this report primarily displayed neurologic signs, similar to those observed in natural H5N1 HPAI infected fox kits (*13*) and a stone marten (*5*) in Europe, and to both naturally and experimentally infected birds of prey (*17,18*). Widespread postmortem lesions and viral detection from multiple tissues in the majority of cases are consistent with systemic viral infections in these mammals. Necrotizing and predominantly non-suppurative meningoencephalitis and acute interstitial pneumonia were the primary microscopic lesions in the mammals in this article, followed by myocardial necrosis and less consistently hepatic necrosis and lymphoid depletion. These findings are consistent with lesions reported in naturally and experimentally infected foxes (*13,19*), domestic cats (*20*), and a stone marten (*5*). A similar distribution of lesions has also been reported in raptors with HPAI, with the brain, heart, and lungs being most affected (*17, 18, 21*). Notably, pancreatic necrosis was often present in birds of prey but was only present in one red fox in this study.

Of note, brain and heart lesions were absent or mild in the striped skunks in this study, with hepatic necrosis, lymphoid necrosis, and interstitial pneumonia being the predominant findings, which correspond with the distribution of viral antigen and nucleic acids as detected by IHC and PCR. A similar distribution of lesions has been reported in domestic cats naturally infected with H5N1 HPAIv (*22*). Therefore, the typical constellation of lesions associated with HPAIv infection cannot be assumed to be uniform across mammalian families.

Immunohistochemical analysis indicated that the virus primarily resides within neuron cell bodies and processes in the brains of infected animals, which corresponded with the brain being the most frequently and strongly positive tissue for PCR detection in most species in this study. Five of 22 animals in which IHC was performed on brain tissue lacked immunoreactivity. In one case (red fox 34), the brain was strongly positive by PCR but only the heart showed immunoreactivity, and the cause of this discrepancy is unclear. In the other four animals that lacked brain immunoreactivity (red fox 12, 15, 19 and bobcat 1), potential causes are discussed below. Conversely, positive IHC reaction within the lungs was rare and scattered in most species, despite the severity of gross and microscopic lung lesions in most cases. The cause of this discrepancy is unclear but may relate to cytokine-induced pulmonary injury, which has been reported in experimental infection trials in laboratory mammals (*23*).

PCR was the most sensitive method for IAV detection in this study, and there were only four cases in which IAV was detected by PCR but tissues lacked immunoreactivity. One red fox with no tissue immunoreactivity (red fox 15) was strongly PCR positive in multiple tissues, and significant tissue autolysis prior to fixation may have interfered with sensitivity of immunohistochemical analysis. In two red foxes, lack of tissue immunoreactivity was likely due to absence or limited amounts of HPAIv in tissue samples from these foxes, which were either weakly positive by PCR (red fox 12) or positive by nasal swab only (red fox 19). The detection of HPAIv in these two foxes may have been clinically incidental, as both had concurrent illness that could have explained their clinical signs and postmortem lesions (*E. coli* septicemia and canine adenovirus, respectively). Similarly, the single grey fox had extensive hemorrhage in multiple body cavities that was interpreted as the cause of mortality and there were no microscopic lesions. This animal was weakly positive on pooled nasal and oropharyngeal swab only, while tracheal swab and brain tissue were IAV negative by PCR. This detection may have represented early or subclinical IAV infection, or simply mucosal carriage of IAV. In one bobcat with brain lesions that lacked immunoreactivity, antigen may have been absent due to chronicity of lesions.

Subclinical infection with HPAIv has been previously documented in experimentally infected 6-to 10-month-old red foxes, which reportedly shed virus in the pharynx for 3 to 7 days in the absence of clinical signs (*19*). In the current study, oropharyngeal swabs were generally more sensitive for the detection of IAV in foxes than were nasal or intestinal/rectal swabs, consistent with higher oropharyngeal viral shedding reported in experimental infections (*19*).

Although fecal and nasal shedding of virus was limited in the experimental infections of 6-to 10-month-old red foxes (*19*), HPAIv was isolated from nasal and oropharyngeal swabs of two red foxes in this study, which presents further evidence that infected red foxes may be capable of transmitting infection.

The majority of clinically ill mammals in this study died or were euthanized due to progression of clinical signs, similar to outcomes reported in other natural infections with H5N1 HPAIv in mammals (*5, 13, 22, 24, 25*). This dataset is unique in that it includes some evidence that clinical resolution is possible in red foxes. Red fox 6 exhibited neurological signs that resolved following supportive care, and it remained clinically normal until being released to the wild. Red fox 1 and its littermate (not tested for IAV at presentation nor included in this dataset) both exhibited neurological signs at presentation, and red fox 1 died despite supportive care. The presumptively HPAIv-positive litter mate of red fox 1 developed blindness but was otherwise clinically normal for two months, after which it developed neurological signs and was euthanized. That fox (the presumptively HPAIv-positive littermate of red fox 1) was diagnosed with toxoplasmosis on postmortem exam and IAV was not detected in postmortem samples.

Adult red foxes, striped skunks, coyotes, and Virginia opossums infected with HPAIv were not identified in this study. Notably, clinical signs were not observed in 6-to 10-month-old foxes that were experimentally infected with HPAIv (*19*). Reasons that clinical infections have been primarily restricted to young mammals are unknown, but could include naive immune systems in juveniles, different exposure risks across age groups, and behavioral differences that may make infected or clinically ill adult mammals less likely to be encountered. Insufficient data are available at present to postulate on risk factors in the small number of affected adult animals, including a fisher, grey fox, raccoon, and two bobcats. It is worth noting that both adult bobcats were co-infected with parvovirus, and concurrent disease may have contributed to the development of clinical disease due to HPAIv infection in these adult animals. Concurrent infections in HPAIv-infected mammals have not been previously reported, but were identified in 5 animals in this study, including parvovirus in bobcats, and one case each of canine adenovirus, rabies, and *E. coli* in red foxes.

Ingestion of birds infected with HPAIv is presumed to be the most likely source of infection in these wild mammals. Wild birds, including waterfowl, are a typical or occasional component of the natural diet in these mammalian species and infection following ingestion of HPAIv-positive birds has been confirmed in red foxes (*19*), domestic dogs (*24*), domestic cats (*25*), captive felids (*26*), and multiple species of raptors and scavenging birds (*17, 18, 21*). Furthermore, for some red foxes in this study, there were reports of vixens bringing dead waterfowl to the den, and three red foxes had evidence of bird ingestion at postmortem examination. Mortalities in HPAIv-infected wild birds and domestic poultry have been reported throughout the United States during this outbreak, and wild birds infected with HPAIv have been confirmed in the majority of counties in which these mammals originated (*3,4*). Wild bird surveillance for HPAIv was widespread in the United States, yet variable from state to state. For example, the lack of HPAIv detections in a given county likely reflects sampling bias, rather than a true absence of virus activity in those locations. Passive and active surveillance efforts have demonstrated the extensive distribution of HPAIv in all migratory bird flyways in North America (*3,4*), further evidence that the virus is widespread geographically. Horizontal transmission of H5N1 HPAIv has been documented in experimentally infected domestic cats (*25*) and ferrets (*26*), and transmission from an infected parent or conspecific cannot be ruled out as a potential source of infection in some of the mammals in this study.

The scattered geographical and temporal distribution of the HPAIv-infected mammals in this article indicates that these infections represent sporadic spillover events into individual animals or litters that are sharing the landscape with HPAIv-infected wild birds. This theory is supported by sequencing data from mammalian samples, confirming the presence of several different genotypes emerging and circulating in the United States in wild birds. Although active surveillance efforts in wild mammals were not initiated following the recognition of these cases, anecdotally there has not been evidence of sustained transmission of HPAIv within wild populations of terrestrial mammalian species in any of the locations represented in this study. In mammals, sustained mortality events due to H5N1 HPAIv have thus far only been reported in seals (*28*), and methods of transmission responsible for the outbreak remain unclear. The E627K substitution that has been associated with mammalian adaptation (*12)* was identified in three mammals in this article, and continued vigilance is warranted as ongoing spillover of avian influenza viruses into mammalian hosts could potentially result in further reassortment or adaptation of these viruses to broader host ranges (*29, 30*).

In conclusion, we demonstrate that multiple North American wild terrestrial mammal species are susceptible to natural infection with H5N1 HPAIv of Eurasian lineage goose/Guangdong H5 2.3.4.4b subtype, likely via ingestion of infected wild birds. Neurological signs were the primary clinical manifestation in these mammals, and HPAIv infection warrants consideration as a differential diagnosis along with more common causes of neurologic disease in wild mammals, such as rabies virus, canine distemper virus, canine adenovirus, and toxoplasmosis. Given the ongoing nature of the current HPAI outbreak, surveillance for HPAIv in wild mammals that share the landscape with or consume wild birds would contribute to a better understanding of the distribution of these viruses in free-ranging wildlife.

## Acknowledgements

The authors wish to acknowledge Logan McCormick and the other staff of Dane County Humane Society’s Wildlife Center, staff of the Wildlife Rehabilitation Center of Minnesota, Tracy Belle of Wildthunder Wildlife & Animal Rehabilitation & Sanctuary, and Erica Zuhlke of Critter Crossings Rehabilitation Center for their astute observations and care of the wild animals included in this article. We are grateful for the expertise, contributions, and collaboration of many wildlife agency experts, including Michelle Carstensen of the Minnesota Department of Natural Resources; Conservation Officer Ben Bergman of the Iowa Department of Natural Resources; Nancy Businga, Carissa Knab, and Mandy Kamps of the Wisconsin Department of Natural Resources; Dr. Stout of the Utah Division of Wildlife Resources; Katie Farinosi, Melinda Cosgrove, Mauri Liberati, Tonya Tepin, and field staff of the Michigan Department of Natural Resources; Sarah Bevins of the National Wildlife Research Center; and the North Dakota Game and Fish Department. Finally, this work would not have been possible without the many laboratory diagnosticians at contributing institutions, including Melissa Fadden of the Cornell University Wildlife Health Lab, Dr. Gauger of the Iowa State University Veterinary Diagnostic Laboratory, Katy Griffin of the U.S. Geological Survey National Wildlife Health Center, and Dr. Kelly and Dr. Hullinger of the Utah Veterinary Diagnostic Laboratory. USGS data can be found at https://doi.org/10.5066/P9KAJA8J.

## Disclaimer

Any use of trade, product or firm names is for descriptive purposes and does not imply endorsement by the U.S. Government. The views expressed in this article are those of the authors and do not necessarily reflect the official policy of the U.S. Department of Agriculture.

