## Supplementary Tables for "Pathology of natural infection with highly pathogenic avian influenza virus (H5N1) clade 2.3.4.4b in wild terrestrial mammals in the United States in 2022"

| **Supplementary Table 1**. Reference sequences used for single nucleotide polymorphism analysis. | | | |
|---|---|---|---|
| Segment | Reference used | GenBank accession | GISAID ID |
| PB2 | A/Blue-Winged Teal/Alberta/39/2020(H4N6) | MZ171362.1 | N/A |
| PB1 | EPI2049863\|A/chicken/NL/FAV-0033/2021 | N/A | EPI_ISL_12968823 |
| PA | EPI2049864\|A/chicken/NL/FAV-0033/2021 | N/A | EPI_ISL_12968823 |
| HA | EPI2049865\|A/chicken/NL/FAV-0033/2021 | N/A | EPI_ISL_12968823 |
| NP | A/mallard/Nevada/AH0190913/2021(H11N6) | OL462886.1 | N/A |
| NA | EPI2049867\|A/chicken/NL/FAV-0033/2021 | N/A | EPI_ISL_12968823 |
| MP | EPI2049868\|A/chicken/NL/FAV-0033/2021 | N/A | EPI_ISL_12968823 |
| NS | EPI2049869\|A/chicken/NL/FAV-0033/2021 | N/A | EPI_ISL_12968823 |

N/A = not applicable

**Supplementary Table 2.** Summary of signalment, date found, days in human care (if applicable), state and county found, and clinical outcome of wild mammals naturally infected with highly pathogenic avian influenza, grouped by species and listed in order of date found.

| Case ID | | Species | Sex | Age class | Date found | Days in care | State | County | Outcome |
|---|---|---|---|---|---|---|---|---|---|
| Red fox | 1 | *Vulpes vulpes* | F | Juv | 4/1/22 | < 1 | MI | Macomb | D |
| Red fox | 2 | *Vulpes vulpes* | M | Juv | 4/1/22 | <1 | NY | Cayuga | E |
| Red fox | 3 | *Vulpes vulpes* | NR | NR | 4/2/22 | NA | AK | Aleutians west census area | D |
| Red fox | 4 | *Vulpes vulpes* | NR | Juv | 4/7/22 | <1 | NY | Erie | E |
| Red fox | 5 | *Vulpes vulpes* | NR | Juv | 4/7/22 | >1 | NY | Erie | E |
| Red fox | 6 | *Vulpes vulpes* | F | Juv | 4/9/22 | >8 | WI | Jefferson | A |
| Red fox | 7 | *Vulpes vulpes* | F | Juv | 4/9/22 | 2 | WI | Jefferson | D |
| Red fox | 8 | *Vulpes vulpes* | M | Juv | 4/9/22 | >1 | NY | Monroe | E |
| Red fox | 9 | *Vulpes vulpes* | F | Juv | 4/11/22 | <1 | NY | Tompkins | E |
| Red fox | 10 | *Vulpes vulpes* | F | Juv | 4/11/22 | <1 | NY | St. Lawrence | E |
| Red fox | 11 | *Vulpes vulpes* | M | Juv | 4/11/22 | >1 | NY | St. Lawrence | E |
| Red fox | 12 | *Vulpes vulpes* | F | Juv | 4/12/22 | 5 | WI | Dane | E |
| Red fox | 13 | *Vulpes vulpes* | M | Juv | 4/14/22 | < 1 | MI | Lapeer | D |
| Red fox | 14 | *Vulpes vulpes* | F | Juv | 4/14/22 | < 1 | MI | St. Clair | D |
| Red fox | 15 | *Vulpes vulpes* | M | Juv | 4/19/22 | < 1 | WI | Rock | E |
| Red fox | 16 | *Vulpes vulpes* | M | Juv | 4/19/22 | <1 | NY | Erie | D |
| Red fox | 17 | *Vulpes vulpes* | NR | Juv | 4/20/22 | NA | MN | Anoka | D |
| Red fox | 18 | *Vulpes vulpes* | M | Juv | 4/24/22 | < 1 | WI | Waushara | E |
| Red fox | 19 | *Vulpes vulpes* | F | Juv | 4/24/22 | < 1 | WI | Dane | E |
| Red fox | 20 | *Vulpes vulpes* | M | Juv | 4/25/22 | <1 | MN | Dakota | E |
| Red fox | 21 | *Vulpes vulpes* | F | Juv | 4/25/22 | <1 | MN | Hennepin | E |
| Red fox | 22 | *Vulpes vulpes* | M | Juv | 5/3/22 | 1 | WI | Adams | E |
| Red fox | 23 | *Vulpes vulpes* | F | Juv | 5/3/22 | 1 | WI | Adams | E |
| Red fox | 24 | *Vulpes vulpes* | F | Juv | 5/4/22 | >8 | WI | Dane | A |
| Red fox | 25 | *Vulpes vulpes* | M | Juv | 5/4/22 | NA | IA | Hancock | E |
| Red fox | 26 | *Vulpes vulpes* | M | Juv | 5/4/22 | <1 | NY | Niagara | E |
| Red fox | 27 | *Vulpes vulpes* | F | Juv | 5/5/22 | 1 | WI | Grant | E |
| Red fox | 28 | *Vulpes vulpes* | M | Juv | 5/5/22 | 1 | WI | Grant | D |
| Red fox | 29 | *Vulpes vulpes* | F | Juv | 5/5/22 | NA | MI | Osceola | E |
| Red fox | 30 | *Vulpes vulpes* | NR | Juv | 5/8/22 | <1 | MN | Otter Tail | E |
| Red fox | 31 | *Vulpes vulpes* | F | Juv | 5/9/22 | 2 | IA | Cedar | E |
| Red fox | 32 | *Vulpes vulpes* | M | Juv | 5/10/22 | NA | MI | Midland | D |
| Red fox | 33 | *Vulpes vulpes* | M | Juv | 5/10/22 | NA | MI | Midland | D |
| Red fox | 34 | *Vulpes vulpes* | F | Juv | 5/10/22 | NA | MN | Stearns | D |
| Red fox | 35 | *Vulpes vulpes* | M | Juv | 5/11/22 | <1 | NY | Allegany | E |
| Red fox | 36 | *Vulpes vulpes* | M | Juv | 5/14/22 | <1 | MI | Gladwin | D |

| | | | | | | | | | |
|---|---|---|---|---|---|---|---|---|---|
| Red fox | 37 | *Vulpes vulpes* | M | Juv | 5/15/22 | <1 | MI | Gladwin | D |
| Red fox | 38 | *Vulpes vulpes* | M | Juv | 5/18/22 | NA | MI | Ostego | D |
| Red fox | 39 | *Vulpes vulpes* | F | Juv | 5/18/22 | <1 | MI | Bay | D |
| Red fox | 40 | *Vulpes vulpes* | M | Juv | 5/19/22 | NA | MI | Osceola | E |
| Red fox | 41 | *Vulpes vulpes* | F | Juv | 5/23/22 | NA | ND | Dickey | E |
| Red fox | 42 | *Vulpes vulpes* | M | Juv | 5/23/22 | NA | ND | Dickey | E |
| Red fox | 43 | *Vulpes vulpes* | M | Juv | 5/24/22 | NA | UT | Salt Lake | E |
| Red fox | 44 | *Vulpes vulpes* | M | Juv | 5/26/22 | NA | UT | Salt Lake | E |
| Red fox | 45 | *Vulpes vulpes* | M | Juv | 5/31/22 | 8 | MI | Mackinac | E |
| Red fox | 46 | *Vulpes vulpes* | M | Juv | 6/1/22 | <1 | MI | Muskegon | D |
| Red fox | 47 | *Vulpes vulpes* | M | Juv | 6/6/22 | NA | AK | Nome census area | E |
| Red fox | 48 | *Vulpes vulpes* | NR | Juv | NR | <1 | MN | NR | E |
| Red fox | 49 | *Vulpes vulpes* | F | Juv | NR | <1 | MN | NR | E |
| Red fox | 50 | *Vulpes vulpes* | M | Juv | NR | <1 | MN | NR | E |
| Skunk | 1 | *Mephitis mephitis* | F | Juv | 6/6/22 | >3 | ID | Latah | D |
| Skunk | 2 | *Mephitis mephitis* | M | Juv | 6/6/22 | >3 | ID | Latah | E |
| Skunk | 3 | *Mephitis mephitis* | F | Juv | 6/7/22 | >3 | ID | Latah | E |
| Skunk | 4 | *Mephitis mephitis* | F | Juv | 6/7/22 | >3 | ID | Latah | E |
| Skunk | 5 | *Mephitis mephitis* | M | Juv | 6/7/22 | >3 | ID | Latah | E |
| Skunk | 6 | *Mephitis mephitis* | NR | Juv | 7/21/22 | NA | ID | Ada | D |
| Raccoon | 1 | *Procyon lotor* | F | Ad | 5/11/22 | NA | MI | Iron | E |
| Raccoon | 2 | *Procyon lotor* | NR | Juv | 6/8/22 | NA | WA | Franklin | D |
| Raccoon | 3 | *Procyon lotor* | NR | Juv | 6/8/22 | NA | WA | Franklin | D |
| Raccoon | 4 | *Procyon lotor* | F | Juv | 6/30/22 | NA | WA | Stevens | NR |
| Bobcat | 1 | *Lynx rufus* | F | Ad | 4/24/22 | NA | WI | Lincoln | E |
| Bobcat | 2 | *Lynx rufus* | F | Ad | 5/6/22 | NA | WI | Lincoln | E |
| Opossum | 1 | *Didelphis virginiana* | M | Juv | 5/18/22 | 1 | IA | Benton | E |
| Opossum | 2 | *Didelphis virginiana* | F | Juv | 5/18/22 | 2 | IA | Benton | D |
| Coyote | 1 | *Canis latrans* | F | Juv | 5/3/22 | NA | MI | Wexford | E |
| Fisher | 1 | *Pekania pennanti* | M | Ad | 4/14/22 | NA | WI | St. Croix | E |
| Grey fox | 1 | *Urocyon cinereoargenteus* | F | Ad | 6/2/22 | NA | MI | Clinton | D |

NR = not recorded; F = female, M = male, Ad = adult, Juv = juvenile, NA = not applicable, A = alive at time of publication, D = died naturally, E = euthanized.

**Supplementary Table 3.** Summary of reported clinical signs demonstrated by wild mammals naturally infected with highly pathogenic avian influenza virus, grouped by species.

| Case ID | | Neurological signs, any reported | Seizures | Ataxia | Tremors | Lack of fear | Hyper-salivation | Lethargy | Febrile | Other, reported in <5 animals |
|---|---|---|---|---|---|---|---|---|---|---|
| Red fox | 1 | X | X | X | | | | | X | |
| Red fox | 2 | X | X | | | | | | | |
| Red fox | 4 | X | X | | | | X | X | X | unconscious |
| Red fox | 5 | X | X | | | | X | X | X | unconscious |
| Red fox | 6 | X | | X | | | | X | | |
| Red fox | 7 | X | | X | | | | X | | vocalization |
| Red fox | 8 | X | | | | | X | | | blindness, circling |
| Red fox | 9 | X | X | | X | | | X | | vocalization |
| Red fox | 10 | X | X | | | | | | | vocalization |
| Red fox | 11 | X | X | | | | | | | diarrhea, vocalization |
| Red fox | 12 | | | | | | | X | | |
| Red fox | 13 | X | X | X | | | | | X | |
| Red fox | 14 | X | X | X | | | | | X | |
| Red fox | 15 | X | X | | X | | | | | vocalization |
| Red fox | 16 | | X | | X | | X | X | | |
| Red fox | 19 | X | | X | | | | | | diarrhea, dyspnea |
| Red fox | 20 | X | X | X | X | | | X | | |
| Red fox | 21 | X | | | X | | | X | | |
| Red fox | 22 | X | X | | X | | | X | | |
| Red fox | 23 | X | | | X | | | X | | torticollis |
| Red fox | 24 | | | | | | | | | dehydrated, thin |
| Red fox | 25 | X | | X | | X | | | | |
| Red fox | 26 | | X | | | | X | X | | |
| Red fox | 27 | X | X | | X | | | X | | nystagmus, torticollis |
| Red fox | 28 | X | X | | | | | | | |
| Red fox | 29 | X | | | | | | | | |
| Red fox | 30 | X | X | | | | | | | blindness, circling |
| Red fox | 31 | X | X | | | | | | | blindness, grimace |
| Red fox | 35 | X | X | | | | | | | dull, vomiting |
| Red fox | 36 | X | | | | | | | | |
| Red fox | 37 | X | | X | | | | X | | recumbent |
| Red fox | 39 | X | | X | | | | X | | |
| Red fox | 40 | X | X | X | | | | | | |
| Red fox | 41 | X | | X | | | | | | unconscious |

| | | | | | | | | | | |
|---|---|---|---|---|---|---|---|---|---|---|
| Red fox | 42 | X | | X | | | | | | paralysis |
| Red fox | 43 | X | X | | | | | | | |
| Red fox | 44 | X | | X | | | | X | | |
| Red fox | 45 | X | | | | | | | | |
| Red fox | 46 | X | | | | X | | X | | |
| Red fox | 47 | X | | | | X | | | | circling |
| Red fox | 48 | X | X | X | X | | | X | | |
| Red fox | 49 | X | X | X | X | | | X | | |
| Red fox | 50 | X | | X | | | | X | | |
| Skunk | 1 | X | | X | X | | | X | | dyspnea |
| Skunk | 2 | X | | X | X | | | X | | dyspnea |
| Skunk | 3 | X | X | | X | | | X | | |
| Skunk | 4 | X | X | | X | | | X | | |
| Skunk | 5 | X | X | | X | | | X | | |
| Raccoon | 1 | X | | | | | | | | |
| Raccoon | 4 | X | | X | | | | | | |
| Bobcat | 1 | X | | | X | X | | | | |
| Bobcat | 2 | X | | | | X | | X | | dyspnea |
| Opossum | 1 | X | X | | | | | | | |
| Opossum | 2 | X | X | | | | | | | |
| Coyote | 1 | X | | X | | | | X | | |
| Fisher | 1 | X | | X | | X | | X | | |
| Grey fox | 1 | X | | X | | X | | | | circling |

X = reported; blank = not reported.

**Supplementary Table 4.** Summary of histopathologic lesion distribution in wild mammals naturally infected with highly pathogenic avian influenza virus infection.

| Case ID | | Brain lesions: Necrotizing and inflammatory meningoencephalitis | | | | | | Myocardial necrosis | Interstitial pneumonia | Lymphoid depletion | Hepatic necrosis |
|---|---|---|---|---|---|---|---|---|---|---|---|
| | | Brainstem | Cerebellum | Hippocampus | Thalamus | Cortex | Frontal lobe | | | | |
| Red fox | 1 | | | | | + | | + | + | + | - |
| Red fox | 2 | + | + | + | | + | | - | + | - | - |
| Red fox | 4 | | | | | | | | | | |
| Red fox | 5 | | | | | | | | | | |
| Red fox | 7 | + | + | + | + | + | | + | + | + | + |
| Red fox | 8 | - | - | - | + | - | + | + | - | - | - |
| Red fox | 9 | + | + | + | + | + | + | + | + | - | - |
| Red fox | 10 | | | | | + | | - | + | - | - |
| Red fox | 12 | - | - | - | - | - | + | + | + | + | + |
| Red fox | 13 | | | | | + | | + | + | + | - |
| Red fox | 14 | | | | | + | | + | + | + | - |
| Red fox | 15 | | | | | | | + | + | + | + |
| Red fox | 17 | + | + | + | + | + | | | + | | |
| Red fox | 18 | + | + | + | + | + | + | + | + | + | - |
| Red fox | 19 | + | | | + | + | + | + | + | + | + |
| Red fox | 20 | + | - | - | + | + | | + | + | | + |
| Red fox | 21 | + | - | + | + | + | | + | + | - | + |
| Red fox | 22 | + | + | - | + | + | + | + | + | - | + |
| Red fox | 23 | + | - | + | + | + | + | + | + | - | + |
| Red fox | 25 | | | | | + | | + | + | + | + |
| Red fox | 26 | + | | + | + | + | + | + | + | - | + |
| Red fox | 27 | + | + | | + | + | + | + | + | - | - |
| Red fox | 28 | + | + | | | + | + | - | + | + | - |
| Red fox | 29 | | | | | + | | + | + | + | + |
| Red fox | 30 | | - | | - | + | | | | | |
| Red fox | 31 | | - | | | + | | + | + | + | + |
| Red fox | 32 | | | | | + | | - | + | A | A |
| Red fox | 33 | | | | | + | | - | + | A | A |
| Red fox | 34 | - | - | + | + | + | | + | + | - | - |
| Red fox | 35 | + | - | | + | + | + | - | + | - | - |
| Red fox | 36 | | | | | + | | - | + | - | - |
| Red fox | 37 | | | | | + | | - | + | A | A |
| Red fox | 38 | | | | | + | | - | - | - | A |
| Red fox | 39 | | | | | + | | - | + | A | A |
| Red fox | 49 | + | + | + | | + | | + | + | - | - |
| Red fox | 41 | | - | + | | + | + | - | + | + | + |
| Red fox | 42 | | - | + | | + | + | + | + | + | - |
| Red fox | 43 | + | + | + | + | + | | + | + | + | - |
| Red fox | 44 | | | - | + | + | | + | - | + | - |
| Red fox | 45 | | | | | + | | - | + | + | - |
| Red fox | 46 | | | | | + | | + | + | + | - |
| Skunk | 1 | + | - | - | - | - | | - | + | + | + |
| Skunk | 2 | + | - | - | - | - | | - | + | + | + |
| Skunk | 3 | - | - | - | - | + | | - | + | + | + |
| Skunk | 4 | - | - | - | - | - | | - | + | + | + |
| Skunk | 5 | - | - | - | - | - | | - | + | + | + |
| Raccoon | 1 | | | | | + | | - | + | - | - |
| Raccoon | 2 | + | - | - | - | + | | - | - | + | - |
| Raccoon | 3 | + | + | - | - | + | | - | + | + | + |
| Raccoon | 4 | - | - | + | + | + | | + | + | - | - |
| Bobcat | 1 | + | - | + | + | + | + | + | + | + | - |
| Bobcat | 2 | + | - | + | + | + | + | + | - | + | + |

| | | | | | | | | | | | |
|---|---|---|---|---|---|---|---|---|---|---|---|
| Opossum | 1 | | | | | + | | - | - | - | + |
| Opossum | 2 | | - | | | - | | - | - | - | - |
| Coyote | 1 | | | | | + | | + | + | - | A |
| Fisher | 1 | | - | + | + | + | + | - | + | + | + |
| Grey fox | 1 | | | | | - | | - | - | - | - |

blank = not evaluated; - = lesion not present; + = lesion present; A = too autolyzed to interpret.

**Supplementary Table 5.** Genotype (where available) and cycle threshold (CT) results for quantitative real time PCR for influenza A virus (IAV), IAV H5 subtype, and avian influenza virus H5 2.3.4.4b subtype assays in wild mammals, by tissue or sample type.

| Case ID | Genotype | Assay | Nasal swab | OP swab | Nasal & OP swab | Tracheal swab | Brain | Lung | Heart | Intestine* | Liver | Kidney | Spleen |
|---|---|---|---|---|---|---|---|---|---|---|---|---|---|
| Red fox 1 | B1.2 | IAV | n/d | n/d | 19.4 | 31.6 | 13.9 | n/d | n/d | n/d | n/d | n/d | n/d |
| | | H5 | n/d | n/d | 17.5 | 28.8 | 12.4 | n/d | n/d | n/d | n/d | n/d | n/d |
| | | 2.3.4.4 | n/d | n/d | 21.5 | 32 | 14.7 | n/d | n/d | n/d | n/d | n/d | n/d |
| Red fox 2 | n/d | IAV | n/d | n/d | n/d | n/d | 15 | n/d | n/d | n/d | n/d | n/d | n/d |
| Red fox 3 | A3 | IAV | 28.4 | n/d | n/d | n/d | n/d | n/d | n/d | n/d | n/d | n/d | n/d |
| | | H5 | 24.5 | n/d | n/d | n/d | n/d | n/d | n/d | n/d | n/d | n/d | n/d |
| | | 2.3.4.4 | 28.2 | n/d | n/d | n/d | n/d | n/d | n/d | n/d | n/d | n/d | n/d |
| Red fox 4 | Minor02 | IAV | n/d | n/d | n/d | n/d | 17.2 | n/d | n/d | n/d | n/d | n/d | n/d |
| | | H5 | n/d | n/d | n/d | n/d | 8.6 | n/d | n/d | n/d | n/d | n/d | n/d |
| | | 2.3.4.4 | n/d | n/d | n/d | n/d | 13.1 | n/d | n/d | n/d | n/d | n/d | n/d |
| Red fox 5 | Minor02 | IAV | n/d | n/d | n/d | n/d | 12.6 | n/d | n/d | n/d | n/d | n/d | n/d |
| | | H5 | n/d | n/d | n/d | n/d | 12.9 | n/d | n/d | n/d | n/d | n/d | n/d |
| | | 2.3.4.4 | n/d | n/d | n/d | n/d | 17.2 | n/d | n/d | n/d | n/d | n/d | n/d |
| Red fox 6 | n/d | IAV | 38.8 | n/d | n/d | n/d | n/d | n/d | n/d | n/d | n/d | n/d | n/d |
| | | H5 | 36 | n/d | n/d | n/d | n/d | n/d | n/d | n/d | n/d | n/d | n/d |
| | | 2.3.4.4 | 38.6 | n/d | n/d | n/d | n/d | n/d | n/d | n/d | n/d | n/d | n/d |
| Red fox 7 | B2 | IAV | 38.9 | 28.5 | 28.3 | 32.1 | 37.8 | 31.3 | n/d | n/d | n/d | n/d | n/d |
| | | H5 | N | 33.6 | 26.2 | 29.6 | N | 29 | n/d | n/d | n/d | n/d | n/d |
| | | 2.3.4.4 | 37.8 | 36.8 | 29 | 31.8 | 39.6 | 31.7 | n/d | n/d | n/d | n/d | n/d |
| Red fox 8 | n/d | IAV | n/d | n/d | n/d | n/d | 38.1 (FFPE) | n/d | n/d | n/d | n/d | n/d | n/d |
| Red fox 9 | n/d | IAV | n/d | n/d | n/d | n/d | n/d | 34.792 | n/d | n/d | n/d | n/d | n/d |
| | | 2.3.4.4 | n/d | n/d | n/d | n/d | n/d | 34.6 | n/d | n/d | n/d | n/d | n/d |
| Red fox 10 | B1.1 | IAV | n/d | n/d | n/d | n/d | 17.104 | n/d | n/d | n/d | n/d | n/d | n/d |
| | | H5 | n/d | n/d | n/d | n/d | 15.4 | n/d | n/d | n/d | n/d | n/d | n/d |
| | | 2.3.4.4 | n/d | n/d | n/d | n/d | 20 | n/d | n/d | n/d | n/d | n/d | n/d |
| Red fox 11 | B1.1 | IAV | n/d | n/d | n/d | n/d | 27.093 | n/d | n/d | n/d | n/d | n/d | n/d |
| | | H5 | n/d | n/d | n/d | n/d | 22 | n/d | n/d | n/d | n/d | n/d | n/d |
| | | 2.3.4.4 | n/d | n/d | n/d | n/d | 25.9 | n/d | n/d | n/d | n/d | n/d | n/d |
| Red fox 12 | n/d | IAV | N | 35.5 | n/d | n/d | N | 38.6 | 40.1 | N | n/d | n/d | n/d |
| | | H5 | n/d | 34 | n/d | n/d | 37.4 | N | N | N | n/d | n/d | n/d |
| | | 2.3.4.4 | n/d | 36 | n/d | n/d | n/d | 39.4 | N | N | n/d | n/d | n/d |
| Red fox 13 | B1.2 | IAV | n/d | n/d | 36.3 | 37.4 | 15.6 | n/d | n/d | N | n/d | n/d | n/d |
| | | H5 | n/d | n/d | 34 | 34.2 | 15.5 | n/d | n/d | n/d | n/d | n/d | n/d |
| | | 2.3.4.4 | n/d | n/d | 38.5 | 37.3 | 17.7 | n/d | n/d | n/d | n/d | n/d | n/d |
| Red fox 14 | B1.2 | IAV | n/d | n/d | 28.6 | 28.6 | 11.9 | n/d | n/d | N | n/d | n/d | n/d |
| | | H5 | n/d | n/d | 25.9 | 26.3 | 10.4 | n/d | n/d | n/d | n/d | n/d | n/d |

| | | | | | | | | | | | | | |
|---|---|---|---|---|---|---|---|---|---|---|---|---|---|
| | | 2.3.4.4 | n/d | n/d | 28.6 | 28.8 | 12.9 | n/d | n/d | n/d | n/d | n/d | n/d |
| Red fox 15 | B2 | IAV | n/d | n/d | n/d | 32.9 | 17.6 | 24.4 | 34.6 | 37.5 | n/d | n/d | n/d |
| | | H5 | n/d | n/d | n/d | 28.2 | 13.9 | 21.3 | 26.2 | 34.8 | n/d | n/d | n/d |
| | | 2.3.4.4 | n/d | n/d | n/d | 32.1 | 16.7 | 24.5 | 28.3 | 36 | n/d | n/d | n/d |
| Red fox 16 | n/d | IAV | n/d | n/d | n/d | n/d | 15 | n/d | n/d | n/d | n/d | n/d | n/d |
| | | N1 | n/d | n/d | n/d | n/d | 14.4 | n/d | n/d | n/d | n/d | n/d | n/d |
| | | 2.3.4.4 | n/d | n/d | n/d | n/d | 16.2 | n/d | n/d | n/d | n/d | n/d | n/d |
| Red fox 17 | B3.2 | IAV | n/d | n/d | n/d | n/d | 13.4 | n/d | n/d | n/d | n/d | n/d | n/d |
| | | H5 | n/d | n/d | n/d | n/d | 14 | n/d | n/d | n/d | n/d | n/d | n/d |
| | | 2.3.4.4 | n/d | n/d | n/d | n/d | 13.7 | n/d | n/d | n/d | n/d | n/d | n/d |
| Red fox 18 | B2 | IAV | 36.9 | 29.1 | n/d | n/d | 29 | 24.5 | 35.4 | 37.5 | n/d | n/d | n/d |
| | | H5 | N | 24.8 | n/d | n/d | 26.6 | 20.9 | 34 | 37.5 | n/d | n/d | n/d |
| | | 2.3.4.4 | 38.3 | 28.4 | n/d | n/d | 29.8 | 23.8 | 36 | 36 | n/d | n/d | n/d |
| Red fox 19 | n/d | IAV | 34.6 | N | n/d | n/d | N | N | N | N | n/d | n/d | n/d |
| | | H5 | 32.9 | n/d | n/d | n/d | n/d | N | N | N | n/d | n/d | n/d |
| | | 2.3.4.4 | 36 | n/d | n/d | n/d | n/d | N | N | N | n/d | n/d | n/d |
| Red fox 20 | B2 | IAV | n/d | n/d | n/d | n/d | 15.9 | n/d | n/d | n/d | n/d | n/d | n/d |
| | | H5 | n/d | n/d | n/d | n/d | 16 | n/d | n/d | n/d | n/d | n/d | n/d |
| | | 2.3.4.4 | n/d | n/d | n/d | n/d | 19 | n/d | n/d | n/d | n/d | n/d | n/d |
| Red fox 21 | B2 | IAV | n/d | n/d | n/d | n/d | 22.4 | n/d | n/d | n/d | n/d | n/d | n/d |
| | | H5 | n/d | n/d | n/d | n/d | 22.1 | n/d | n/d | n/d | n/d | n/d | n/d |
| | | 2.3.4.4 | n/d | n/d | n/d | n/d | 24.7 | n/d | n/d | n/d | n/d | n/d | n/d |
| Red fox 22 | Minor15 | IAV | 29.8 | 23.8 | n/d | n/d | 13.5 | 32.3 | 33.3 | 34.5 | n/d | n/d | n/d |
| | | H5 | 29.1 | 22.2 | n/d | n/d | 14.6 | 30.7 | 33.7 | 33.8 | n/d | n/d | n/d |
| | | 2.3.4.4 | 28.7 | 24.9 | n/d | n/d | 15 | 30.9 | 33.2 | 33.5 | n/d | n/d | n/d |
| Red fox 23 | Minor15 | IAV | 39.8 | 32.9 | n/d | n/d | 16.4 | 34 | 33.5 | N | n/d | n/d | n/d |
| | | H5 | N | 30.7 | n/d | n/d | 16.5 | 32.1 | 32.8 | 38.1 | n/d | n/d | n/d |
| | | 2.3.4.4 | 36 | 33.6 | n/d | n/d | 16.4 | 32.5 | 32.4 | 37.9 | n/d | n/d | n/d |
| Red fox 24 | n/d | IAV | N | 36 | n/d | n/d | n/d | n/d | n/d | n/d | n/d | n/d | n/d |
| | | H5 | N | 36.9 | n/d | n/d | n/d | n/d | n/d | n/d | n/d | n/d | n/d |
| | | 2.3.4.4 | N | 35.9 | n/d | n/d | n/d | n/d | n/d | n/d | n/d | n/d | n/d |
| Red fox 25 | B3.2 | IAV | n/d | 25.2 | 26.9 | n/d | 23.4 | n/d | n/d | N | n/d | n/d | n/d |
| | | H5 | n/d | 27.2 | 29 | n/d | 26 | n/d | n/d | N | n/d | n/d | n/d |
| | | 2.3.4.4 | n/d | 27.4 | 28.5 | n/d | N | n/d | n/d | n/d | n/d | n/d | n/d |
| Red fox 26 | n/d | IAV | n/d | n/d | n/d | n/d | n/d | 32.3 | n/d | n/d | n/d | n/d | n/d |
| | | H5 | n/d | n/d | n/d | n/d | n/d | 32 | n/d | n/d | n/d | n/d | n/d |
| | | 2.3.4.4 | n/d | n/d | n/d | n/d | n/d | 33.3 | n/d | n/d | n/d | n/d | n/d |
| Red fox 27 | B3.1 | IAV | 25.8 | 32.2 | n/d | n/d | 12.9 | 24.8 | 24.7 | 30.9 | n/d | n/d | n/d |
| | | H5 | 24.2 | 30.6 | n/d | n/d | 11.6 | 24.3 | 22.9 | 31.4 | n/d | n/d | n/d |
| | | 2.3.4.4 | 27.6 | 35.1 | n/d | n/d | 14.8 | 28.2 | 28 | 34.3 | n/d | n/d | n/d |
| | B3.1 | IAV | 21.3 | 23.2 | n/d | n/d | 12.9 | 21.1 | 21.3 | 30.3 | n/d | n/d | n/d |

| | | | | | | | | | | | | | |
|---|---|---|---|---|---|---|---|---|---|---|---|---|---|
| Red fox 28 | | H5 | 18.7 | 21.5 | n/d | n/d | 11.1 | 19.9 | 19 | 29.7 | n/d | n/d | n/d |
| | | 2.3.4.4 | 22.8 | 25.7 | n/d | n/d | 14.7 | 23.7 | 23.6 | 33.5 | n/d | n/d | n/d |
| Red fox 29 | B1.2 | IAV | 29.8 | n/d | n/d | n/d | n/d | n/d | n/d | n/d | n/d | n/d | n/d |
| | | H5 | 32.3 | n/d | n/d | n/d | n/d | n/d | n/d | n/d | n/d | n/d | n/d |
| | | 2.3.4.4 | 34.1 | n/d | n/d | n/d | n/d | n/d | n/d | n/d | n/d | n/d | n/d |
| Red fox 30 | B3.2 | IAV | n/d | n/d | n/d | n/d | 19.8 | n/d | n/d | n/d | n/d | n/d | n/d |
| | | H5 | n/d | n/d | n/d | n/d | 19.8 | n/d | n/d | n/d | n/d | n/d | n/d |
| | | 2.3.4.4 | n/d | n/d | n/d | n/d | 19.9 | n/d | n/d | n/d | n/d | n/d | n/d |
| Red fox 31 | B1.2 | IAV | n/d | N | 34.9 | n/d | 29.8 | n/d | n/d | N | n/d | n/d | n/d |
| | | H5 | n/d | n/d | 38.3 | n/d | 33 | n/d | n/d | n/d | n/d | n/d | n/d |
| | | 2.3.4.4 | n/d | n/d | N | n/d | 30.9 | n/d | n/d | n/d | n/d | n/d | n/d |
| Red fox 32 | B1.1 | IAV | 26.4 | n/d | n/d | n/d | n/d | n/d | n/d | n/d | n/d | n/d | n/d |
| | | H5 | 28.2 | n/d | n/d | n/d | n/d | n/d | n/d | n/d | n/d | n/d | n/d |
| | | 2.3.4.4 | 29 | n/d | n/d | n/d | n/d | n/d | n/d | n/d | n/d | n/d | n/d |
| Red fox 33 | B2 | IAV | 32.9 | n/d | n/d | n/d | n/d | n/d | n/d | n/d | n/d | n/d | n/d |
| | | H5 | 35.1 | n/d | n/d | n/d | n/d | n/d | n/d | n/d | n/d | n/d | n/d |
| | | 2.3.4.4 | 36.3 | n/d | n/d | n/d | n/d | n/d | n/d | n/d | n/d | n/d | n/d |
| Red fox 34 | n/d | IAV | n/d | N | n/d | n/d | 31.5 | n/d | n/d | n/d | n/d | n/d | n/d |
| | | H5 | n/d | n/d | n/d | n/d | 29.1 | n/d | n/d | n/d | n/d | n/d | n/d |
| | | 2.3.4.4 | n/d | n/d | n/d | n/d | 30.5 | n/d | n/d | n/d | n/d | n/d | n/d |
| Red fox 35 | n/d | IAV | n/d | n/d | n/d | n/d | n/d | 39.2 | n/d | n/d | n/d | n/d | n/d |
| | | H5 | n/d | n/d | n/d | n/d | n/d | 39.4 | n/d | n/d | n/d | n/d | n/d |
| | | 2.3.4.4 | n/d | n/d | n/d | n/d | n/d | N | n/d | n/d | n/d | n/d | n/d |
| Red fox 36 | B1.1 | IAV | 29.2 | 23.6 | n/d | n/d | 14.3 | n/d | n/d | n/d | n/d | n/d | n/d |
| | | H5 | 26.4 | 26.1 | n/d | n/d | 15.7 | n/d | n/d | n/d | n/d | n/d | n/d |
| | | 2.3.4.4 | 30 | 26 | n/d | n/d | 16.8 | n/d | n/d | n/d | n/d | n/d | n/d |
| Red fox 37 | B1.1 | IAV | 39.6 | 22.7 | n/d | n/d | n/d | n/d | n/d | n/d | n/d | n/d | n/d |
| | | H5 | 37.6 | 25.4 | n/d | n/d | n/d | n/d | n/d | n/d | n/d | n/d | n/d |
| | | 2.3.4.4 | 38.9 | 25.1 | n/d | n/d | n/d | n/d | n/d | n/d | n/d | n/d | n/d |
| Red fox 38 | B1.2 | IAV | 27 | n/d | n/d | n/d | n/d | n/d | n/d | n/d | n/d | n/d | n/d |
| | | H5 | 27 | n/d | n/d | n/d | n/d | n/d | n/d | n/d | n/d | n/d | n/d |
| | | 2.3.4.4 | 27.3 | n/d | n/d | n/d | n/d | n/d | n/d | n/d | n/d | n/d | n/d |
| Red fox 39 | B1.1 | IAV | 33.3 | 25 | n/d | n/d | 18.1 | n/d | n/d | n/d | n/d | n/d | n/d |
| | | H5 | 30.1 | 24.8 | n/d | n/d | 18.4 | n/d | n/d | n/d | n/d | n/d | n/d |
| | | 2.3.4.4 | 32.6 | 34.9 | n/d | n/d | 19.6 | n/d | n/d | n/d | n/d | n/d | n/d |
| Red fox 40 | n/d | IAV | 36.3 | n/d | n/d | n/d | n/d | n/d | n/d | n/d | n/d | n/d | n/d |
| | | H5 | 36.8 | n/d | n/d | n/d | n/d | n/d | n/d | n/d | n/d | n/d | n/d |
| | | 2.3.4.4 | 37 | n/d | n/d | n/d | n/d | n/d | n/d | n/d | n/d | n/d | n/d |
| Red fox 41 | B3.1 | IAV | n/d | 27.93 | n/d | n/d | n/d | n/d | n/d | n/d | n/d | n/d | n/d |
| | | H5 | n/d | 31.4 | n/d | n/d | n/d | n/d | n/d | n/d | n/d | n/d | n/d |
| | | 2.3.4.4 | n/d | 30.57 | n/d | n/d | n/d | n/d | n/d | n/d | n/d | n/d | n/d |

| | | | | | | | | | | | | | |
|---|---|---|---|---|---|---|---|---|---|---|---|---|---|
| Red fox 42 | B3.1 | IAV | n/d | 27.61 | n/d | n/d | n/d | n/d | n/d | n/d | n/d | n/d | n/d |
| | | H5 | n/d | 30.28 | n/d | n/d | n/d | n/d | n/d | n/d | n/d | n/d | n/d |
| | | 2.3.4.4 | n/d | 30.55 | n/d | n/d | n/d | n/d | n/d | n/d | n/d | n/d | n/d |
| Red fox 43 | B4 | IAV | n/d | n/d | n/d | N | 27.5 | n/d | n/d | n/d | n/d | n/d | n/d |
| | | H5 | n/d | n/d | n/d | N | 28 | n/d | n/d | n/d | n/d | n/d | n/d |
| | | 2.3.4.4 | n/d | n/d | n/d | n/d | 29 | n/d | n/d | n/d | n/d | n/d | n/d |
| Red fox 44 | n/d | IAV | n/d | n/d | n/d | N | 29.9 | n/d | n/d | n/d | n/d | n/d | n/d |
| | | H5 | n/d | n/d | n/d | n/d | N | n/d | n/d | n/d | n/d | n/d | n/d |
| | | 2.3.4.4 | n/d | n/d | n/d | n/d | 33.8 | n/d | n/d | n/d | n/d | n/d | n/d |
| Red fox 45 | n/d | IAV | 32.7 | n/d | n/d | n/d | 38.7 | n/d | n/d | n/d | n/d | n/d | n/d |
| | | H5 | 34.4 | n/d | n/d | n/d | N | n/d | n/d | n/d | n/d | n/d | n/d |
| | | 2.3.4.4 | 39.5 | n/d | n/d | n/d | 39 | n/d | n/d | n/d | n/d | n/d | n/d |
| Red fox 45 | B3.2 | IAV | 24.9 | 26.4 | n/d | n/d | 20.5 | n/d | n/d | n/d | n/d | n/d | n/d |
| | | H5 | 21.7 | 27.6 | n/d | n/d | 22.6 | n/d | n/d | n/d | n/d | n/d | n/d |
| | | 2.3.4.4 | 26.4 | 28.4 | n/d | n/d | 24.1 | n/d | n/d | n/d | n/d | n/d | n/d |
| Red fox 46 | B3.2 | IAV | 32.9 | n/d | n/d | n/d | n/d | 26.4 | n/d | 33.4 | n/d | n/d | n/d |
| | | H5 | 30.4 | n/d | n/d | n/d | n/d | 24.7 | n/d | 32.0 | n/d | n/d | n/d |
| | | 2.3.4.4 | 31.2 | n/d | n/d | n/d | n/d | 26.2 | n/d | 33.0 | n/d | n/d | n/d |
| Red fox 47 | B2 | IAV | n/d | n/d | 24.3 | n/d | n/d | n/d | n/d | n/d | n/d | n/d | n/d |
| | | H5 | n/d | n/d | 24.1 | n/d | n/d | n/d | n/d | n/d | n/d | n/d | n/d |
| | | 2.3.4.4 | n/d | n/d | 27.2 | n/d | n/d | n/d | n/d | n/d | n/d | n/d | n/d |
| Red fox 48 | B3.2 | IAV | n/d | n/d | 33.7 | n/d | n/d | n/d | n/d | n/d | n/d | n/d | n/d |
| | | H5 | n/d | n/d | 32.1 | n/d | n/d | n/d | n/d | n/d | n/d | n/d | n/d |
| | | 2.3.4.4 | n/d | n/d | 34.5 | n/d | n/d | n/d | n/d | n/d | n/d | n/d | n/d |
| Red fox 49 | n/d | IAV | n/d | n/d | 23.9 | n/d | n/d | n/d | n/d | n/d | n/d | n/d | n/d |
| | | 2.3.4.4 | n/d | n/d | 23.9 | n/d | n/d | n/d | n/d | n/d | n/d | n/d | n/d |
| Skunk 2 | B3.2 | IAV | n/d | n/d | n/d | n/d | 14.5 | 20.5 | n/d | n/d | n/d | n/d | n/d |
| | | H5 | n/d | n/d | n/d | n/d | 11.1 | 19 | n/d | n/d | n/d | n/d | n/d |
| | | 2.3.4.4 | n/d | n/d | n/d | n/d | 11.3 | 18.5 | n/d | n/d | n/d | n/d | n/d |
| Skunk 3 | B3.2 | IAV | n/d | n/d | n/d | n/d | n/d | 19.1 | n/d | n/d | 19.4‡ | n/d | n/d |
| | | H5 | n/d | n/d | n/d | n/d | n/d | 18.5 | n/d | n/d | 19.6‡ | n/d | n/d |
| | | 2.3.4.4 | n/d | n/d | n/d | n/d | n/d | 18.2 | n/d | n/d | 20.1‡ | n/d | n/d |
| Skunk 4 | B3.2 | IAV | n/d | n/d | n/d | n/d | 22 | 19.2 | n/d | n/d | 20.4‡ | n/d | n/d |
| | | H5 | n/d | n/d | n/d | n/d | 20.7 | 19 | n/d | n/d | 20.5‡ | n/d | n/d |
| | | 2.3.4.4 | n/d | n/d | n/d | n/d | 23.6 | 20 | n/d | n/d | 20.8‡ | n/d | n/d |
| Skunk 6 | n/d | IAV | n/d | 28.7 | n/d | n/d | 20.9 | n/d | n/d | n/d | n/d | n/d | n/d |
| | | H5 | n/d | N | n/d | n/d | N | n/d | n/d | n/d | n/d | n/d | n/d |
| | | 2.3.4.4 | n/d | 27.3 | n/d | n/d | 19.9 | n/d | n/d | n/d | n/d | n/d | n/d |
| Raccoon 1 | B1.2 | IAV | 35.6 | 39.9 | n/d | n/d | 23.1 | n/d | n/d | n/d | n/d | n/d | n/d |
| | | H5 | 36.5 | N | n/d | n/d | 23.9 | n/d | n/d | n/d | n/d | n/d | n/d |

| | | | | | | | | | | | | | |
|---|---|---|---|---|---|---|---|---|---|---|---|---|---|
| | | 2.3.4.4 | 34.3 | 39.6 | n/d | n/d | 25 | n/d | n/d | n/d | n/d | n/d | n/d |
| Raccoon 2 | B2 | IAV | n/d | 32.1 | n/d | n/d | n/d | n/d | n/d | n/d | n/d | n/d | n/d |
| | | H5 | n/d | 29.2 | n/d | n/d | n/d | n/d | n/d | n/d | n/d | n/d | n/d |
| | | 2.3.4.4 | n/d | 32.9 | n/d | n/d | n/d | n/d | n/d | n/d | n/d | n/d | n/d |
| Raccoon 4 | n/d | IAV | n/d | 34.5 | n/d | n/d | n/d | n/d | n/d | n/d | n/d | n/d | n/d |
| | | H5 | n/d | 31.6 | n/d | n/d | n/d | n/d | n/d | n/d | n/d | n/d | n/d |
| | | 2.3.4.4 | n/d | 34 | n/d | n/d | n/d | n/d | n/d | n/d | n/d | n/d | n/d |
| Opossum 1 | B1.2 | IAV | n/d | 23.1 | 23.4 | n/d | 17.5 | n/d | n/d | N | n/d | n/d | n/d |
| | | H5 | n/d | 25 | 25.1 | n/d | n/d | n/d | n/d | n/d | n/d | n/d | n/d |
| | | 2.3.4.4 | n/d | 24.8 | 24.7 | n/d | 24.9 | n/d | n/d | n/d | n/d | n/d | n/d |
| Opossum 2 | B1.2 | IAV | n/d | 30.9 | 28.1 | n/d | 16.2 | n/d | n/d | N | n/d | n/d | n/d |
| | | H5 | n/d | 35 | 32.4 | n/d | n/d | n/d | n/d | n/d | n/d | n/d | n/d |
| | | 2.3.4.4 | n/d | 36 | 33.2 | n/d | 23.9 | n/d | n/d | n/d | n/d | n/d | n/d |
| Bobcat 1 | n/d | IAV | 34.3 | 35.2 | n/d | n/d | 20.4 | 27.6 | 38.8 | 34.7 | n/d | n/d | n/d |
| | | H5 | 29.8 | 30.5 | n/d | n/d | 20.5 | 28.5 | N | 36 | n/d | n/d | n/d |
| | | 2.3.4.4 | 33.9 | 35 | n/d | n/d | 19.1 | 27.2 | 39.1 | 34 | n/d | n/d | n/d |
| Bobcat 2 | B3.2 | IAV | 33.1 | 31.9 | n/d | n/d | 30.3 | N | N | n/d | n/d | n/d | n/d |
| | | H5 | 28.7 | 27.3 | n/d | n/d | 31.3 | 39.6 | N | 34.9 | n/d | n/d | n/d |
| | | 2.3.4.4 | 32.8 | 31.6 | n/d | n/d | 29.9 | N | N | n/d | n/d | n/d | n/d |
| Coyote 1 | B1.2 | IAV | n/d | n/d | 28 | 29.3 | n/d | n/d | n/d | n/d | n/d | n/d | n/d |
| | | H5 | n/d | n/d | 29 | 27.6 | n/d | n/d | n/d | n/d | n/d | n/d | n/d |
| | | 2.3.4.4 | n/d | n/d | 29.6 | 30.9 | n/d | n/d | n/d | n/d | n/d | n/d | n/d |
| Grey fox 1 | n/d | IAV | n/d | n/d | 37.7 | N | N | n/d | n/d | n/d | n/d | n/d | n/d |
| | | H5 | n/d | n/d | 37.4 | N | N | n/d | n/d | n/d | n/d | n/d | n/d |
| | | 2.3.4.4 | n/d | n/d | 35.7 | N | N | n/d | n/d | n/d | n/d | n/d | n/d |
| Fisher 1 | B3.2 | IAV | n/d | 29.7 | n/d | n/d | 17 | 22.2 | n/d | 33.9 | 33 | 28.7 | 31 |
| | | H5 | n/d | 28.6 | n/d | n/d | 19.4 | 24.5 | n/d | 34.8 | 34.9 | 30.5 | 32.8 |
| | | 2.3.4.4 | n/d | 28.6 | n/d | n/d | 19.3 | 24.8 | n/d | 35.4 | 34.4 | 30.2 | 31.5 |

OP = oropharyngeal; * = includes rectal swabs, intestinal swabs, and intestine sample types; N = not detected; n/d = not done; ‡ = pooled liver and spleen.

**Supplementary Table 6**. Summary of results for select viral etiologies in wild mammals naturally infected with highly pathogenic avian influenza. All testing performed is by real time PCR, unless otherwise indicated.

| Case ID | | HPAI | AIV^ | RV* | CAV | CDV | CHV | CPV/FPV | CCoV | CRCoV | CPIV | FHV | FCV | FCoV | SARS-CoV-2 |
|---|---|---|---|---|---|---|---|---|---|---|---|---|---|---|---|
| Red fox | 1 | P | | N | | N^ | | | | | | | | | |
| Red fox | 2 | P | | N | | N^ | | | | | | | | | |
| Red fox | 3 | P | | | | | | | | | | | | | |
| Red fox | 4 | P | | | | | | | | | | | | | |
| Red fox | 5 | P | | | | | | | | | | | | | |
| Red fox | 6 | P | | | | N | | | | | | | | | |
| Red fox | 7 | P | P | N | N | N | N | N | N | N | N | | | | |
| Red fox | 8 | P | | N | | N^ | | | | | | | | | |
| Red fox | 9 | P | P | N | | | | | | | | | | | |
| Red fox | 10 | P | | N | | | | | | | | | | | |
| Red fox | 11 | P | | N | | N | | | | | | | | | |
| Red fox | 12 | P | N | N | N | N | N | N | N | N | N | | | | |
| Red fox | 13 | P | | N | | N^ | | | | | | | | | |
| Red fox | 14 | P | | N | | N^ | | | | | | | | | |
| Red fox | 15 | P | N | N | N | N | N | N | N | N | N | | | | |
| Red fox | 16 | P | | | | | | | | | | | | | |
| Red fox | 17 | P | P | N | | N^ | | | | | | | | | |
| Red fox | 18 | P | P | | N | N | N | N | N | N | N | | | | |
| Red fox | 19 | P | N | N | P | N | N | N | N | N | N | | | | |
| Red fox | 20 | P | P | N | | N^ | | | | | | | | | |
| Red fox | 21 | P | P | N | | N^ | | | | | | | | | |
| Red fox | 22 | P | P | N | N | N | N | N | N | N | N | | | | |
| Red fox | 23 | P | P | N | N | N | N | N | N | N | N | | | | |
| Red fox | 24 | P | | | | | | | | | | | | | |
| Red fox | 25 | P | P | N | | N | | | | | | | | | |
| Red fox | 26 | P | | N | | | | | | | | | | | |
| Red fox | 27 | P | P | | N | N | N | N | N | N | N | | | | |
| Red fox | 28 | P | P | | N | N | N | N | N | N | N | | | | |
| Red fox | 29 | P | P | | | | | | | | | | | | |
| Red fox | 30 | P | | N | | N^ | | | | | | | | | |
| Red fox | 31 | P | P | N | | N | | | | | | | | | |
| Red fox | 32 | P | | | | | | | | | | | | | |
| Red fox | 33 | P | | | | | | | | | | | | | |
| Red fox | 34 | P | P | N | | N^ | | | | | | | | | |
| Red fox | 35 | P | | N | | | | | | | | | | | |
| Red fox | 36 | P | | N | | | | | | | | | | | |
| Red fox | 37 | P | | N | | | | | | | | | | | |

| | | | | | | | | | | | | | | | |
|---|---|---|---|---|---|---|---|---|---|---|---|---|---|---|---|
| Red fox | 38 | P | | | | | | | | | | | | | |
| Red fox | 39 | P | | N | | | | | | | | | | | |
| Red fox | 40 | P | | | | | | | | | | | | | |
| Red fox | 41 | P | | | | N | | | | | | | | | |
| Red fox | 42 | P | | | | N | | | | | | | | | |
| Red fox | 43 | P | | N | | | | | | | | | | | |
| Red fox | 44 | P | | N | | | | | | | | | | | |
| Red fox | 45 | P | | N | | | | | | | | | | | |
| Red fox | 46 | P | | N | | | | | | | | | | | |
| Red fox | 47 | P | | P | | N | | | | | | | | | |
| Red fox | 48 | P | | | | | | | | | | | | | |
| Red fox | 49 | P | | | | | | | | | | | | | |
| Red fox | 50 | P | | | | | | | | | | | | | |
| Skunk | 1 | | P | | N | N | | | | | | | | | |
| Skunk | 2 | P | P | | | | | | | | | | | | |
| Skunk | 3 | P | P | | | | | | | | | | | | |
| Skunk | 4 | P | P | | | | | | | | | | | | |
| Skunk | 5 | | P | | | | | | | | | | | | |
| Skunk | 6 | P | | N(PCR) | | N^ | | | | | | | | | |
| Raccoon | 1 | P | | N | | | | | | | | | | | |
| Raccoon | 2 | P | P | | | | | | | | | | | | |
| Raccoon | 3 | | P | | | | | | | | | | | | |
| Raccoon | 4 | P | P | | | | | | | | | | | | |
| Bobcat | 1 | P | N | | | N | | P | | | | N | N | N | N |
| Bobcat | 2 | P | P | | | N | | P | | | | N | N | N | N |
| Opossum | 1 | P | P | N | | | | | | | | | | | |
| Opossum | 2 | P | P | N | | | | | | | | | | | |
| Coyote | 1 | P | | | | | | | | | | | | | |
| Fisher | 1 | P | | | | N | | N | | | | | | | N |
| Grey fox | 1 | P | | N | | | | | | | | | | | |

HPAI = highly pathogenic avian influenza virus; AIV = avian influenza virus; RV = rabies virus; CAV = canine adenovirus; CDV = canine distemper virus; CHV = canine herpesvirus; CPV/FPL = canine parvovirus/feline panleukopenia virus; CCoV = canine coronavirus type 1; CRCoV = canine respiratory coronavirus; CPIV = canine parainfluenza virus; FHV = feline herpesvirus; FCV = feline calicivirus; FCOV = feline coronavirus; SARS-CoV-2 = severe acute respiratory syndrome coronavirus 2 (COVID-19); P = positive result; N = negative result; blank = not performed; * = fluorescent antibody testing; ^ = immunohistochemical assay.
